# Individualized AI-driven neuromodulation enhances tongue motor and sensory control networks: a precision intervention for neurorehabilitation in cancer survivors and patients with neurodegenerative disease

**DOI:** 10.1101/2025.04.30.651533

**Authors:** T.D. Papageorgiou, J. Webb, A. Allam, S. Reddy, D. Huynh, R. Hekmati, E.M. Rohren, E.M. Sturgis, S.G. Hilsenbeck, K.A. Hutcheson, S.R. Heilbronner, A.M. Thompson, C.R. Neblett, E Froudarakis

## Abstract

Precise modulation of brain networks responsible for tongue motor and sensory control (TMSC) is critical for restoring functions, such as speech and swallowing in neurodegenerative disease or in treatment-induced chronic cranial neuropathy. We present an individualized, AI-driven fMRI neuromodulation (iNM) platform that adaptively targets subject-specific TMSC networks in real time. To enhance iNM precision and encodability —critical for neurorehabilitation—we mapped each healthy participant’s individualized TMSC selectivity network, creating a subject-specific TMSC digital twin. iNM increased signal strength, spatial expansion, and consistency across motor, sensory, and attention regions, while it reduced signal variability. The bilateral inferior parietal lobule emerged as key sensorimotor integration hub, as it exhibited exclusive activation under iNM along with highest discriminability, and largest spatial expansion. iNM also significantly strengthened and expanded motor, sensory, and attention-related networks — medial-middle frontal areas, insula-claustrum, S1, M1, basal ganglia, motor cerebellum, and inferior temporal— supporting interoceptive and proprioceptive-motor integration. Machine learning and unsupervised hidden Markov modeling revealed that iNM enhanced the decodability and stability of TMSC-neural states, while it suppressed competing swallow-neural state interference. Notably, the iNM effects extended beyond the neuromodulation window, indicating functional persistence—a key requirement for rehabilitation. iNM reconfigured TMSC networks by strengthening cortico-subcortical connectivity and adaptive circuit dynamics. Our findings show iNM as a non-invasive, personalized intervention capable of selectively enhancing sensorimotor control with high spatiotemporal specificity. By demonstrating mechanistic network-precision and functional carryover, iNM offers a promising intervention for individuals with limited treatment options, including head and neck cancer survivors and early-stage neurodegenerative disease patients.

## Introduction

Interoception and proprioception are fundamental processes by which the nervous system monitors internal signals and orchestrates movement. Interoception governs awareness of internal states such as pressure, touch, and pain, while proprioception encodes muscle position, movement, and force. Both are essential for regulating complex orofacial functions, such as tongue motor and sensory control (TMSC)—which underpins swallowing, eating, and speech. These processes are mediated by cranial nerves IX, X, and XII, which receive direct input from primary sensory and motor cortices.

Disruption of TMSC following cranial nerve IX, X, and XII damage is a defining feature of chronic cranial neuropathy (CCN), often arising from surgery or radiotherapy for head and neck cancers.^1–20^ CCN impairs speech, swallowing, and mastication through progressive atrophy and fibrosis, substantially diminishing quality of life.^19^ Despite the increased incidence in cases annually with 10% of survivors developing CCN,^21–23^ the available pharmacologic agents are poorly tolerated and result in adverse drug interactions.^23,24^ TMSC dysfunction also presents early in neurodegenerative conditions, such as amyotrophic lateral sclerosis (ALS), where up to 80% of patients develop dysarthria and dysphagia, culminating in respiratory failure and premature death.^25,26^ In both CCN and ALS, tongue weakness is accompanied by cortical disorganization—including sensory–motor “smudging”—reflecting degraded network precision and excitability. Two lines of evidence support targeting cortical TMSC regions: first, CCN directly compromises tongue elevation, closure, and base retraction; and second, motor cortex stimulation has shown promise in modulating pain, albeit with limitations.^1–3,27–38^ Surface electrical stimulation faces anatomical and physiological barriers, including inconsistent access to deep muscle targets and paradoxical activation of antagonistic pathways. These challenges highlight the need for non-invasive, circuit-specific neuromodulatory strategies to restore tongue control with precision and safety.^4,39–43^

Recent advances in precision neurotechnology have merged high-resolution neural interfaces — such as noninvasive digital holographic imaging^44^ and invasive AI-driven digital twins of the brain in animal models^45,46^ — enabling a long-term goal of patient-specific intervention in real time. These interfaces not only capture neural activity with unprecedented spatial and temporal resolution but also embody digital twins that replicate brain network dynamics and allow real-time prediction of individual responses to therapeutic interventions. This convergence bridges neuroscience and clinical practice, promising tailored neurotherapies that could replace trial-and-error approaches with spatiotemporally precise interventions for conditions ranging from neurological to psychiatric disorders.

In close resemblance to digital twin models of the brain, our real-time, fMRI-based, AI-driven, closed-loop neuromodulation system encapsulates a high-fidelity, individualized representation of a brain network continuously updated using neurophysiological data to reflect the participant’s dynamic brain-level responses to a specific function. Unlike traditional simulations, this neurofunctional twin does not simulate hypothetical neural activity; instead, it is grounded in each individual’s functional network, whose spatially precise and temporally resolved activity is used to dynamically guide iNM-based modulation and optimize performance:^47^ (1) It can target precisely each patient’s unique cortical and subcortical networks involved in tongue motor and sensory control (TMSC), with millimeter-level spatial resolution—capabilities beyond traditional neuromodulation tools like transcranial magnetic stimulation; (2) It is individualized at the anatomical and functional levels, allowing modulation to be adapted to individual neural organization and physiology (**Figure 2**); (3) It operates in a spatiotemporally guided, closed-loop fashion, identifying time-resolved signal peaks or troughs in real time and reinforces (e.g., e.g., enhances motor activity) or suppresses (e.g., decreases pain activity) specific neural patterns to optimize the physiological response under investigation; (4) It is adaptive, continuously adjusting to each patient’s evolving TMSC brain dynamics, with the goal of restoring balanced bilateral activation, resembling healthy network function (**Figure 1**). Thus, iNM can be used as a measurement and a modulation platform and provide guidance based on each participant’s unique neural architecture during TMSC. Notably, success is not defined by BOLD amplitude alone, but also by the adaptive adjustment and spatial expansion of the individualized network’s functional response. As the individualized network is repeatedly engaged through the TMSC physiological response. iNM increases activation thresholds, making maximal activation harder to reach and thereby encouraging more efficient and distributed recruitment.

**Figure 1.**
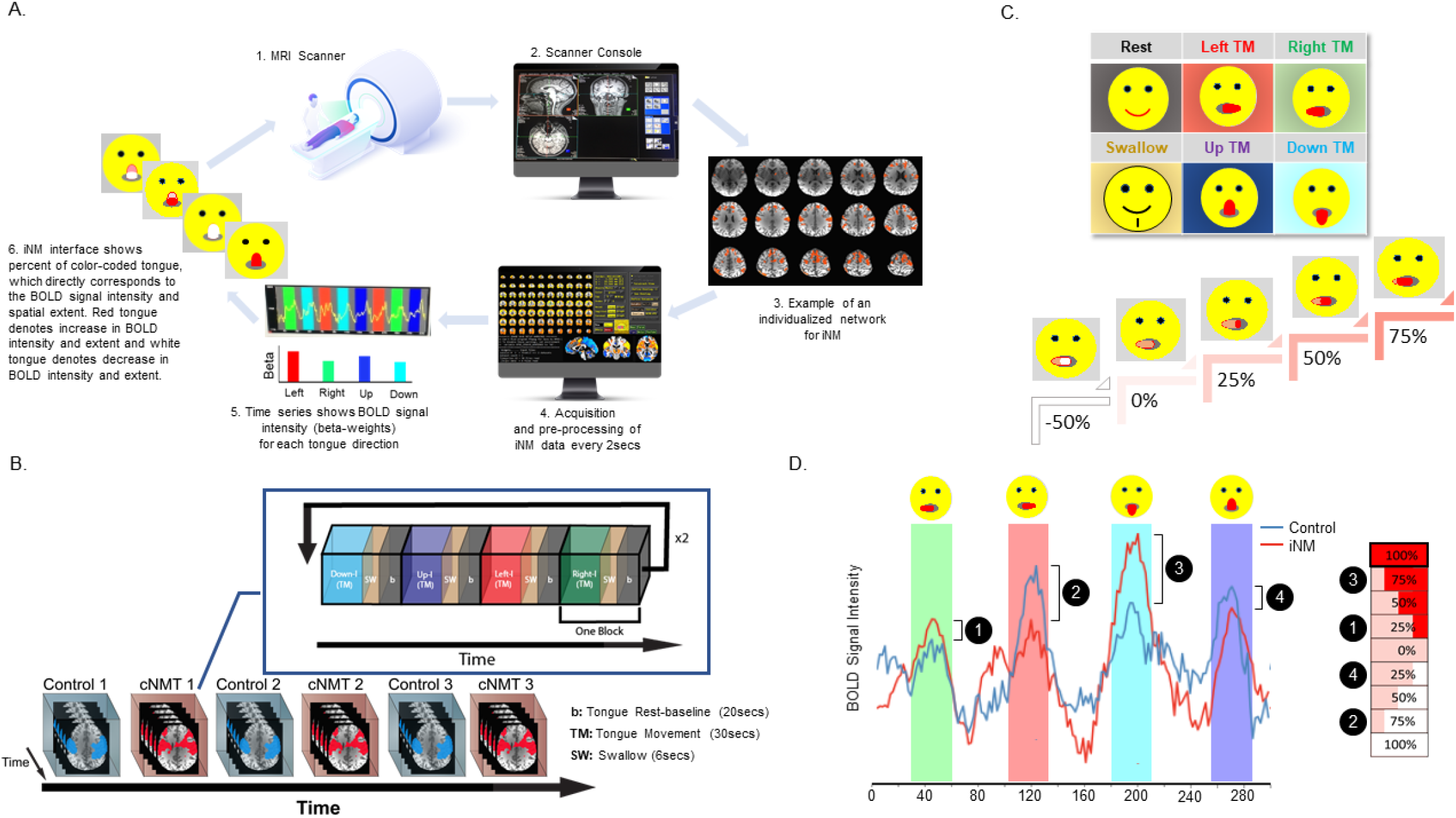
Individualized AI-driven neuromodulation (iNM) **A**. 1. and 2. MRI acquisition of high resolution anatomical images, registered to the Siemens’ console computer; 3. delineate and contour each participant’s individualized motor and sensory network that corresponds to tongue motor and sensory control (TMSC) selectivity, as a function of the upregulated BOLD intensity and spatial extent that corresponds to each condition – tongue movement direction (left; right; up; down); 4. in real-time every TR = 2 seconds, pre-process the iNM data extracted from the individualized network and apply a general linear model to decode each TMSC direction (yellow line = time series, baseline - tongue at rest interval is represented with gray and TMSC red, green, blue, cyan corresponds to left, right, up, and down direction selectivity, respectively); 5. compute the BOLD signal intensity and spatial extent (individualized network) for each direction via a GLM, and the beta-weights are updated as described under **D**; 6. iNM interface shows the extent of color-coded tongue, which directly corresponds to the percent upregulation (red fill-in of the tongue) or downregulation (white fill-in of the tongue) of the BOLD signal intensity and spatial extent. **B**. The temporal sequence of the physiological responses that participants were asked to engage in were the same for the control-No iNM and iNM scans: each TMSC task direction block was 30 seconds long, interleaved with 6-second-duration blocks of swallow and 20-second-duration blocks dedicated to baseline-tongue-at-rest. The order of tongue directions was counterbalanced across runs, and blocks cuing the same tongue direction were not stacked back-to-back. **C**. iNM interface. The entire tongue was filled-in with red during control runs to motivate participants to maximally upregulate the targeted network for each tongue direction. During the iNM scans, the tongue color corresponded to the following scenarios. The BOLD magnitude and extent presents the subject’s individualized brain network that regulates his/her TMSC physiology, which is presented to the participant via the emoji’s tongue: 100%, 75%, 50%, 25% upregulation denoted in red fill-in of the tongue; 0% baseline denoted in pink fill-in of the tongue; −25%, −50%, −75%, −100% downregulation denoted in white fill-in of the tongue. **D**. Computation of the iNM signal. Percent change of the BOLD signal intensity during TMSC compared to baseline-tongue at rest during each control run served as a reference (10th to 90th percentile; shaded in blue) for computing the iNM signal. If the percent BOLD signal change during a 2-second interval was higher than the 10th percentile, then the iNM interface indicated a 10% upregulation, and if it was less than the 10th percentile, the iNM interface indicated downregulation.

**Figure 2.**
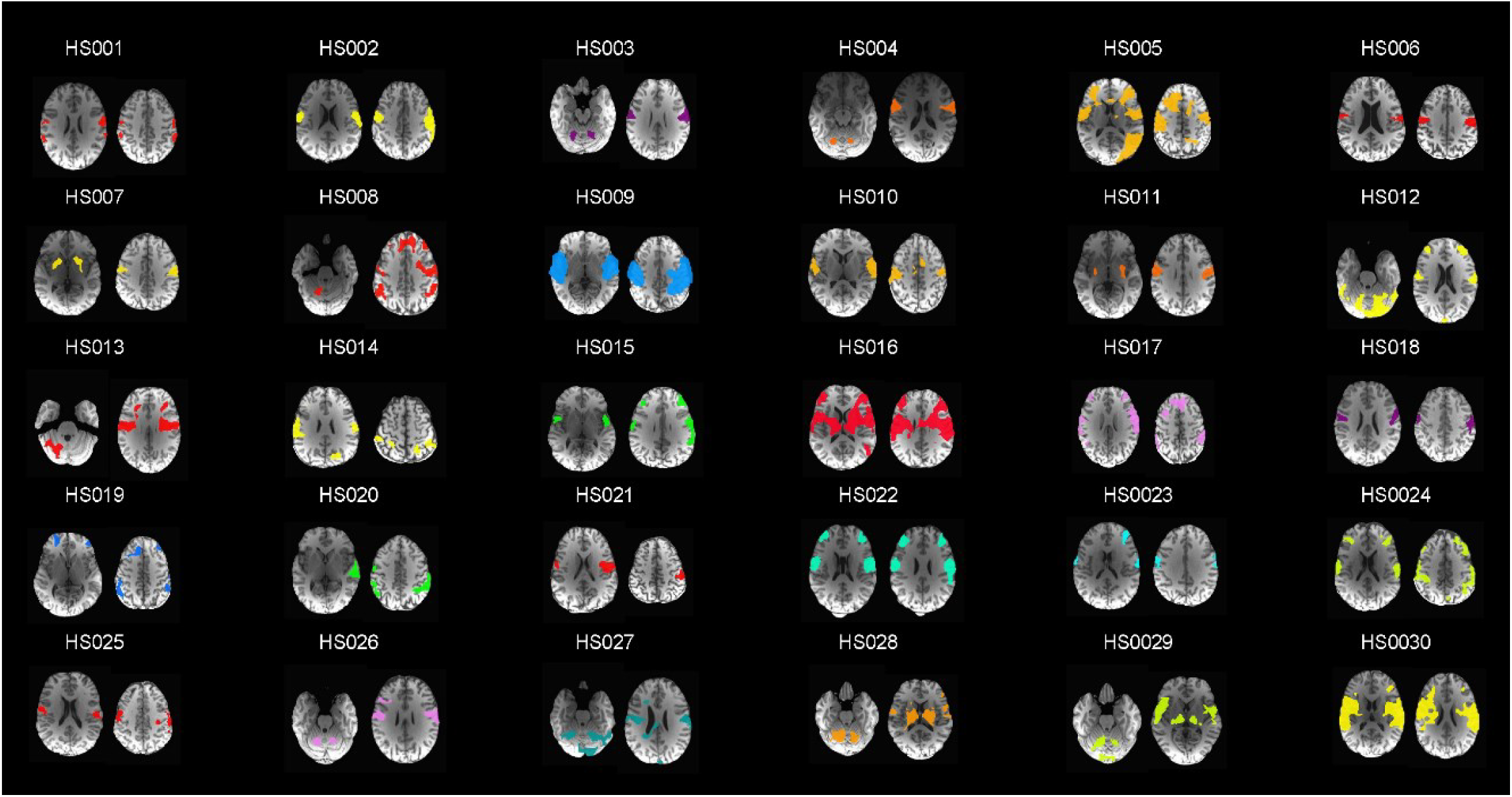
Individualized networks for iNM (n=30). Optimization and efficacy of iNM is enhanced by carefully delineating the intensity and extent of the cortical network based on the patient’s unique anatomy while the TIMSC physiological response is registered. Localization of TMSC is performed by delineating and contouring of each patient’s motor, sensory networks associated with maximal upregulation of tongue movement across directions. Prior to iNM, we found the intersection between the anatomical region and the functional intensity of each subject, while engaged in tongue movement. We then optimized each participant’s network by carefully eliminating areas that represent noise. For example, BOLD activity extending into the cerebrospinal fluid, failing to span multiple slices, or lacking spatial expansion was excluded from analysis.

To realize the potential of individualized, adaptive neuromodulation, it is crucial an intervention like iNM to demonstrate that targeted modulation can reliably and selectively engage specific brain networks. In this study, we focus on TMSC networks—critical for functions, such as swallowing, mastication and speech—as a testbed for assessing the feasibility and network-level mechanisms of iNM. Establishing robust, directionally specific modulation within these networks is a foundational step toward future applications in clinical populations. Specifically, the main objective of this study was to investigate the potential for modulating the activity of motor and sensory networks associated with the tongue-interoceptive and -proprioceptive responses through iNM.

## Materials and Methods Subjects

Thirty healthy, right-handed volunteers (19 males, 11 females, age range = 25.37±5.56) were recruited in this two-day study after informed consent was given in accordance with the Institutional Research Board of Baylor College of Medicine. Exclusion criteria included prior and current medical or psychiatric diagnoses, intake of any medications except over the counter, and general contraindications to MRI examinations. Participants had no history of head, neck, or brain radiation or surgery, no neurological deficits, and normal or corrected-to-normal vision using MR-compatible glasses. At the end of each day, subjects received compensation for volunteering their time.

## MRI and fMRI Pulse Sequence Parameters

Structural and functional brain imaging was performed at the Core for Advanced Magnetic Resonance Imaging (aka CAMRI), Baylor College of Medicine, Houston, Texas using a 3.0 T Siemens Prisma (Siemens, Erlangen, Germany). We used a 20-channel head/neck receiver-array coil for image acquisition. T1-weighted 3D magnetization-prepared gradient-echo (MPRAGE) sequence acquired 192 high-resolution axial slices (field-of-view (FOV) = 245×245 mm2; base resolution = 256×256; repetition time (TR) = 1,200ms; echo time (TE) = 2.66ms; flip angle (FA) = 12°). Functional data consisted of 33 interleaved axial slices, which were acquired using an Echo Planar Imaging (EPI) sequence (FOV = 200×200mm2, voxel size = 3.1×3.1×3.0 mm, TR = 2000ms; flip angle = 90°, number of volumes = 244).

## Real-time fMRI Neuromodulation Acquisition

Turbo-BrainVoyager (TBV; 2.0; Brain Innovation, Maastricht, The Netherlands)^112^ software performed the following preprocessing computations on the EPI images acquired every tongue repetition (TR): (i) 3D motion correction; (ii) incremental linear detrending to remove BOLD signal drifts; (iii) display of statistical brain maps generated from a general linear model (GLM) along with beta weights (BOLD signal intensity values) for each condition; (iv) extraction of the average BOLD signal intensity values from the individualized networks acquired on Day 1 scans (see *Data Analysis*); and (v) presentation of the network’s average BOLD signal intensity via the neuromodulation interface (Figure 1D). The steps of the iNM intervention are summarized in **Figure 1**. To further increase our signal-to-noise ratio (SNR), we employed an exponential moving average (EMA) algorithm to high-pass filter the ROI’s BOLD average and suppress low-frequency components of noise, such as scanner drifts, and physiological noise effects, such as heart rate and respiration. The EMA output of was then low pass filtered via a Kalman filter to eliminate high frequency noise.^113^ Details on how the EMA and Kalman filter were computed can be found in Allam and colleagues (2024).^47^

### Study Structure

Our study was comprised of two day-sessions, separated by 5-7 days. Day 1 consisted of four functional (EPI) scans: three functional task scans to delineate the iNM network and one iNM scans. Day 2 consisted of six EPI scans: three control-no iNM scans, which alternated with three iNM scans. Each EPI scan was composed of 8 continuous periods that lasted for a total of 8 minutes and 12 seconds. Within each period, the subjects were cued to move their tongue in one of four directions (up; down; left; right). Each tongue movement block lasted 30secs and was interleaved with a swallow block (6 secs) and a baseline-tongue at rest block (20 secs). Direction of the movement blocks were randomly counterbalanced across scans and followed three rules: (i) each TMSC direction block occurred twice during each period; (ii) a TMSC direction block was never followed by the same TMSC direction cue; (iii) each scan consisted of a unique order of blocks. **(Figure 1B)**

All subjects were instructed by the principle investigator (TDP) to foster consistency in the subjects’ receipt of instructions and training. These standardized instructions minimized confounding factors unrelated to tongue movement and promoted consistent TMSC, enabling reliable replication across subsequent scans. Subjects were coached to move their tongue without engaging other facial or jaw movements prior to participating in the study. After brief coaching, all 30 subjects were able to perform isolated tongue movements and were included in the study. Study participants were asked to control movement of their tongue in a consistent manner: i. force of tongue movement, and ii. location of tongue’s placement in the mouth.

During the 30-sec tongue movement period, subjects were also coached to abstain from clenching their teeth, pursing their lips together, squinching their eyes, applying pressure to their cheeks, straining their facial muscles, such as those of their forehead between the eyebrows and corners of the eyes, and moving their lower jaw from any backward, forward and side to side movement. Subjects were instructed to move and place their tongue in the same location in their mouth by mentally assigning landmarks, such as a specific ridge on the roof of their palate or, a specific tooth for left and right movement of the tongue. A time interval was dedicated to swallow after each TMSC block. We could detect in real-time if a subject was swallowing during the TMSC task as the fMRI signal would be significantly upregulated during swallowing compared to TMSC. Subjects adhered to abstinence from swallowing during the TMSC task execution and thus, we did not have to exclude swallowing from the TMSC-dedicated time series signal. During the rest block, which served as the baseline, participants stabilized their tongue between the hard and soft palate

### Study Design

#### Overview

On Day 1, we acquired three control-no-iNM scans to delineate each participant’s individualized network. These networks captured each participant’s unique TMSC BOLD amplitude and spatial extent during tongue movement as determined by general linear model (GLM). During the control scans, the visual cue (tongue of the emoji; **Figures 1A, 1C**) was color-coded in red to motivate subjects to execute the task at maximal performance. Each individualized network served as the targets during iNM. iNM was applied in real-time every two seconds during the tongue movement blocks.

#### Computing Individualized ROIs for Neuromodulation Delivery

We acquired functional localizer scans on Day 1 to identify each participant’s individualized network that controls tongue motor and sensory control. After scanning, the acquired anatomical and functional images were processed offline using AFNI.^114^ The anatomical data was spatially transformed to Talairach space using an average volume of 452 skull-stripped brains (TT_icbm452). The functional data was preprocessed to reduce artifacts and increase SNR. Our signal pre-processing protocol included: 1. removal of outliers (head motion, physiological artifacts; 3dDespike) from the time series; 2. slice-time correction (3dTShift); 3. transformation of our oblique-acquired functional dataset to a cardinal dataset (3dWarp -oblique2card); 4. motion correction by registering each functional dataset to the first volume of the first functional scan, using a 3D-rigid-body transformation; 5. spatial smoothing with a 6 mm-FWHM Gaussian kernel filter; and 6. co-registration of the functional data with the individual T1-weighted 3D-structrual data.

The three control-no-iNM scans acquired on Day 1 were then concatenated to increase the SNR prior to generating parametric brain maps across conditions via a General Linear Model (GLM; 3dREMLfit to adjust for temporal correlations in each voxel’s time series, AFNI). The GLM included: 1. four regressors, one for each of the four TMSC directions (left; right; up; down cortical direction selectivity); 2. six covariate vectors that controlled for head motion; 3. white matter and cerebrospinal fluid means were regressed out to increase our SNR, since activity in these areas represents noise; and iv. the baseline-tongue at rest between the hard and soft palate. Each of the four regressors corresponded to each TMSC condition, which was convolved with a gamma variate function, a canonical hemodynamic response function. We excluded the swallow condition from the GLM, since we did not control for the number and onset of swallows performed. We removed the first 3 TRs (6 seconds) from our baseline-tongue-at-rest condition to account for the lag of the hemodynamic response. Following the generation of a network of regions for TMSC compared to baseline-tongue-at-rest (voxelwise p-value=0.001 or 0.005), a cluster analysis was performed to identify highly significant regions (FWER<0.05; **Figure 2)**.

#### Neuromodulation Paradigm

These individualized networks -- each participant’s unique BOLD amplitude and spatial extent – served as the iNM targets with the goal to achieve maximum upregulation during tongue movement **(Figure 1A)**. The color and extent of the emoji’s tongue, which directly represented the magnitude and spatial extent of each subject’s individualized network, conveyed the degree of the network’s upregulation or downregulation in response to tongue movement **(Figure 1C)**. The iNM signal was calculated by first acquiring the percent BOLD signal change (% BOLD PSC) generated during each control scan for each TMSC direction compared to the rest block that preceded it. We computed the % BOLD PSC change from each participant’s individualized areas every 2 seconds as follows:

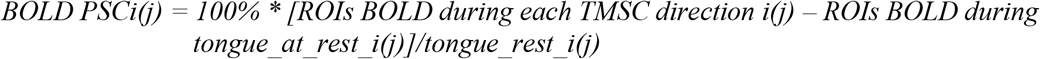

where *i* represents each TMSC direction (left, right, up, down), *j* represents the time interval (2secs) from which the BOLD PSC of tongue motor control direction *i* was computed. iNM presented at each TR (2 seconds) was computed by comparing the current PSC value with a PSC reference range that included 7 bins of 25% BOLD increments: -100%; -75%; -50%; - 25%; 0; 25%; 50%; 75%; 100% **(Figure 1E)**. During the iNM scan that followed each control scan, if the PSC at a given time point was within the reference range or higher than the maximum value, then the tongue was filled-in with red, which represented upregulation of the BOLD signal of the targeted network that controlled a specific tongue movement **(Figure 1E)**. If the PSC though was lower than the minimal value then, the tongue was filled-in with white, and represented downregulation of the BOLD signal of the targeted network that controlled a specific tongue movement **(Figure 1E)**.

#### Support Vector Machine Classification for Direction of Controlled and Voluntary Tongue Movement

Linear support vector machines (SVM) were used to decode BOLD spatial patterns of each TMSC direction generated by the 3 iNM and 3 control conditions. The 3dSVM AFNI function implements a linear SVM classification where the weight vector specifies the signal direction for classification in the multivariate image space.^115–117^

We applied a cross-validation method for each cortical selectivity classification: 1. concatenated all TMSC directions; 2. left direction; 3. right direction; 4. up direction; 5. down direction vs. baseline-tongue at rest. For each subject, we computed classification accuracies (CAs) for the following SVM training-testing permutations by obtaining the means from a 3-fold cross-validation method: i. training and testing with control data and ii. training and testing with iNM data **(Figure SI-1)**. We used soft margin SVMs to account for data points not linearly separable and since we found no differences in performance over a wide range of c - regularization parameter - values from 10-4 to 10+4 (geometric progression with a ratio of 10), we applied the default c=1 for all classifications to optimize training and testing errors.

SVM brain maps were generated for each TMSC direction and condition (control or iNM) as follows: a. the weight vectors corresponding to the optimal hyperplanes obtained from training each of the 3 cross-validation datasets were mapped and mean-zero standardized; b. the weight vectors generated for each subject were normalized; c. the average weight vectors across our 30 subjects were subjected into a t-test for group and cluster analyses (3dttest++ -ClustSim, AFNI); and d. t-test generated brain maps were z-scored (p= 0.005, and cluster size > 10 voxels). From the activation map for each TMSC direction and condition (control or iNM), we generated each cluster’s local maxima (3dExtrema, AFNI). Whether more than one local maximum was generated from a cluster or, a cluster was contained within an anatomical region, only the voxels with the highest z scores which corresponded to increased BOLD signal intensity were included for that region.

#### Percent Signal Change (PSC) Area Under the Curve

We computed the BOLD percent signal change (PSC) for each TMSC direction and condition as a function of time. ROIs were categorized into motor, sensory, attention, and reward networks. To generate the BOLD PSC time series, we first averaged the voxel-wise time series within each ROI, then averaged across segments corresponding to blocks of the same tongue motor control direction from scans of the same type. Each segment began at the onset of a TMSC block and ended at the final time point of the swallow block. For each subject, with 3 control and 3 iNM scans on Day 2, and each scan containing 2 TM blocks per direction, the PSC time series for each direction, condition, and subject was calculated as the average over 6 BOLD signal segments. To compute the AUC, we first averaged the PSC across blocks for each time point, subject, and ROI (resulting in a time × subject × ROI matrix). Next, we averaged these values across subjects to yield a time × ROI matrix, and then across ROIs to produce a single time series. Finally, the Simpson’s method was applied to this time series to estimate the average AUC for the network. The mean area under the PSC curve of each condition (control or iNM) was calculated using Simpson’s Rule:

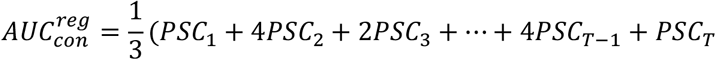

Where PSCt is the PSC at the corresponding time t.

The BOLD percent signal change was then calculated using the following formula:

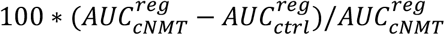

#### D’ Sensitivity Index

To measure the strength of iNM over time and rank regions with respect to this strength, we computed the D’ sensitivity index, also known as Cohen’s d. This index quantifies the separation between the means of the control and iNM distributions, relative to their standard deviations. Assume that the fMRI data set has *T* time points (29 secs), *B* blocks (6 blocks), *S* subjects (n=30), and *N* ROIs, the number of which range between 32-36 dependent on each direction (left direction 32 ROIs; right 36 ROIs; up direction 35; down direction 32). Each ROI is characterized by a by a three-dimensional matrix of dimensions *T*(29) × *B*(6) × *S*(30). Each subject then is characterized by a two-dimensional matrix of dimensions *T* (29) × *B* (6). For a fixed time point *t*, let the distribution of blocks for control and iNM be denoted as 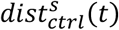 and 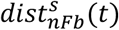, respectively. D’ for the subjects at time point *t* is:

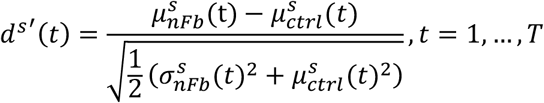

Where 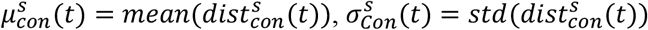.

The sensitivity index between the two conditions at time point *t* and for fixed region is the average sensitivity index across subjects:

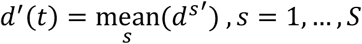

For each subject, bootstrapped distributions of *d*′ time series across all ROIs were divided into bins corresponding to the 5th, 15th, 25th, …, 95th percentile thresholds. At each time point, the *d*′ value for a given ROI was assigned a discretized score from 1 to 10, reflecting the percentile bin in which it resided. The average *d*′ score for each network was then, determined by calculating the mean of the *d*′ scores from all ROIs within that network.

Additionally, peak times were estimated by first applying cubic interpolation to each region’s *d*′ score to generate a smoothed time series. The earliest peak was then identified using MATLAB’s “findpeaks” function with default parameters. These peak times are visualized in the heatmap representation **(Figures 5B, 5C)**.

#### Network-Specific Mapping of Spatial Activation Extent

ROIs were first identified using a general linear model (GLM) thresholded at p=0.001, and FDR-corrected, q ≤ 0.01. Surviving clusters were then assigned to one of four non-overlapping functional networks: motor, sensory, attention, and reward. This classification was guided by anatomical criteria from the Talairach brain atlas and TMSC-specific functional activation patterns. For each subject and each region, we computed the number of activated voxels to quantify the spatial extent of activation, as shown in **Figures 5A; 5D; 5E**. We then calculated the proportion of suprathreshold voxels within each anatomically defined ROI relative to that region’s total template-space volume, yielding a normalized activity index that accounts for differences in region size. These proportions were converted to z-scores across subjects, which provided standardized activation values for cross-subject and network-level comparisons. This approach enabled quantification and comparison of functional network engagement while controlling for anatomical constraints and inter-subject variability.

#### Autoregressive Hidden Markov Model (ARHMM)

To elucidate the dynamic latent network states underlying TMSC during tongue movement—and to validate findings from BOLD signal quantification using the sensitivity index (d′), and AUC—we applied an unsupervised ARHMM. A step-by-step overview of ARHMM analysis is provided in **Figure SI-2**. Briefly, the BOLD data were organized by stacking trials into a single dimension to enable ARHMM processing. The ARHMM was trained in an unsupervised manner on BOLD time series from TMSC-selective regions during tongue movement (6-32 seconds), learning late state dynamics and transition probabilities directly from the data. Inference via the Viterbi trace was applied to the entire time series (0-56 seconds), characterizing brain states during TMSC, swallow, and baseline intervals for each condition **(Figures 6A, SI-9--12)**.

Initially, a single ARHMM was fit to both conditions combined, after which the frequency of latent states was calculated separately for iNM and control trials. For each of the four TMSC directions under iNM and control conditions, we concatenated the sixTMSC blocks across subjects to generate a dataset of dimensions *N × M × T* (ROIs × Trials × Time points), where *M = B × S* reflects the total number of blocks (*B = 6*) across subjects (*S = 30*). After collapsing across all four directions, this yielded a dataset of size *N × 4M × T*, comprised of 720 trials per condition. We performed a leave-one-subject-out analysis, applying the ARHMM to data from 29 subjects (696 trials) while holding out 24 trials from one subject in each iteration, resulting in 30 models per condition. The resulting autoregressive (AR) coefficients generated average ROI-to-ROI connectivity matrices **(Figures SI-13—16)**, which were grouped by functional networks (motor, sensory, attention, reward). Network-level connectivity matrices were computed, and within each network, the summed differences in AR coefficients between control and iNM conditions quantified iNM-induced connectivity changes.

Connectivity measures within and between the Motor, Sensory, Attention, and Reward networks were calculated as the AR coefficients between regions comprising each network.

## Results

The immediate goals of this study were to demonstrate the feasibility of iNM in its guided regulation of proprioceptive and interoceptive control, specifically, targeting TMSC areas and understand its spatiotemporal causal mechanisms. The key to modulating the magnitude and extent successfully is based on the delineation of each patient’s individualized anatomy and physiological dynamics that regulate TMSC, prior to the intervention (**Figure 2)**. We hypothesized that iNM will induce greater consistency in brain TMSC-associated patterns, as well as greater signal-to-noise ratio (SNR), and reduced variance in the magnitude of the signal. To elucidate the causal mechanisms of iNM, we quantified our findings by performing machine learning, advanced computational modeling, and dynamic causal modeling.

### iNM enhances the reliability of BOLD signal intensity and spatial extent

To elucidate the mechanisms of iNM, we first quantified how well we can predict the direction of tongue movement based on the fMRI activity using linear support vector machine (SVM) analysis. SVM-generated cortical maps (**Figure SI-1**) demonstrate enhanced signal intensity and spatial recruitment under iNM, reflecting greater directional selectivity during tongue movement tasks (**Figures 3 and SI-3**). This enhancement in signal intensity and spatial recruitment and area under the receiver operating characteristic curve (AUC-ROC), indicates greater consistency and reliability of the underlying neural signal under iNM (**Figure SI-3**). For all tongue movements combined versus baseline, iNM demonstrated significantly higher mean classification accuracy and AUC-ROC (94.3%, 0.997) compared to control (89.6%, 0.985; *p* < 0.0001). Similarly, for the up and down directions versus baseline, iNM yielded greater accuracy and AUC-ROC values (Up: 95.9%, 0.959; Down: 96.5%, 0.964) relative to control (Up: 90.0%, 0.991; Down: 90.8%, 0.995; *p* < 0.0001).

**Figure 3.**
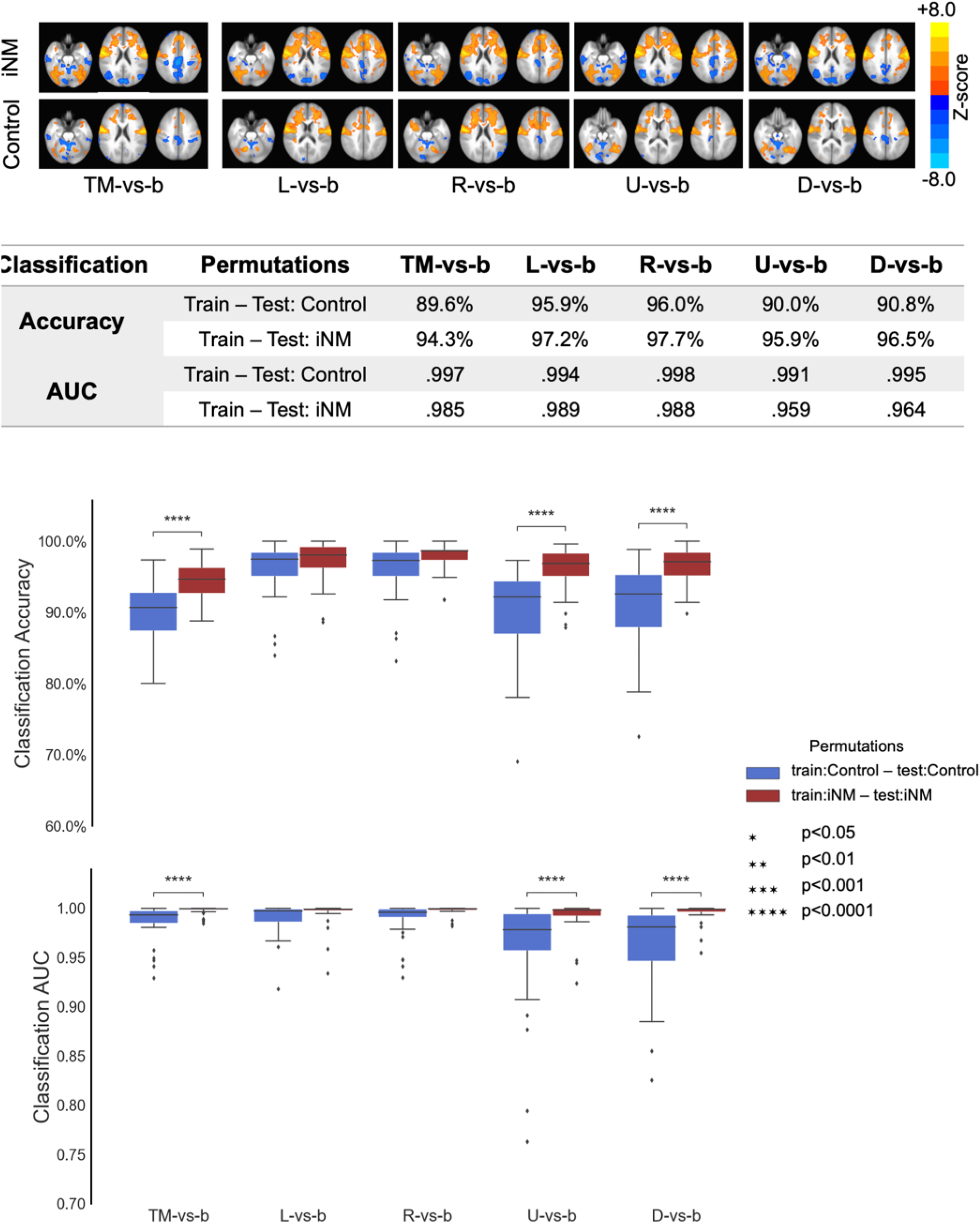
iNM increases consistency in the signal intensity and spatial extent compared to the control-no iNM condition using linear support vector machine analysis. **A**. Decoding cortical networks that control selectivity for each tongue motor control direction. SVM-generated activation map for each tongue movement direction in iNM (top) and control – no iNM (bottom) conditions: voxelwise p value = 0.005, cluster size > 10 voxels. **B**. Table of SVM-generated classification accuracies and areas under the receiver-operator curve. **C**. Boxplot diagram of SVM-generated classification accuracies and areas under the receiver-operator curve (CA, y-axis) across all subjects as a function of different classifications (x-axis) and permutations of training-testing data types (color-coded). L=Left movement, R=Right movement, U=Up movement, D=Down movement, TM=Tongue Movement in all directions, b=Tongue Rest-baseline. On each box, the central line, the lower and upper edges represent the median (Q2), the 25th (Q1) and 75th (Q3) percentiles, respectively. The black open dots represent outliers (bigger than Q3 by 1.5*(Q3-Q1) or smaller than Q1 by 1.5*(Q3-Q1)). The whiskers extend to the most extreme non-outlier datapoints. The black lines and asterisks show significant differences as indicated by paired-samples Wilcoxon sign tests.

#### iNM improves BOLD signal magnitude and reduces its overall variance, as reflected by increased discriminability (d′) and AUC of network-specific ROIs across TMSC conditions over time (**Figure 4**)

The percent change in AUC reflects the modulatory effect of iNM on signal magnitude during each TMSC interval. In the *motor* network, iNM increased mean AUC and d’ scores by 21.7% and 61.6%, respectively across networks with direction-specific AUC changes of 17% (left), 20% (right), 24% (up), 23% (down), and a total BOLD magnitude increase of 847% (**Figures 4A, 5A, 5B, SI-6--8)** and reduced the AUC change in BOLD intensity variance by 9% across TMSC directions **(Figures 4C, SI-8B)**. In the *sensory* network, iNM increased mean AUC by 46% and d’ scores by 60% across networks with direction-specific AUC changes of 41% (left), 44.5% (right), 45% (up), 51.6% (down), and a total BOLD magnitude increase of 2560% (**Figures 4A, 5A, 5B, SI-6--8**). iNM reduced the AUC change in BOLD intensity variance by 14% across TMSC directions **(Figures 4, SI-8B)**. In the *attention* network, iNM increased mean AUC by 37% overall and d’ score of 59% across networks with direction-specific AUC increases of 28% (left), 40% (right), 49% (up), 40% (down), and a total BOLD magnitude increase of 900% (**Figures 4, 5A, 5B, SI-6--8**). iNM reduced the AUC change in BOLD intensity variance by 22% across TMSC directions **(Figures 4, SI-8B)**. In the *reward* network, iNM increased mean AUC by 30%, and a d’ score of 63.3% across networks with direction-specific AUC changes of 7.5% (left), 40.9% (right), 45.2% (up), 32.7% (down), and a total BOLD magnitude increase of 387% (**Figures 4, 5A, 5B, SI-6--8**), while it increased the AUC change in BOLD intensity variance by 3% across TMSC directions **(Figures 4, SI-8B)**.

**Figure 4.**
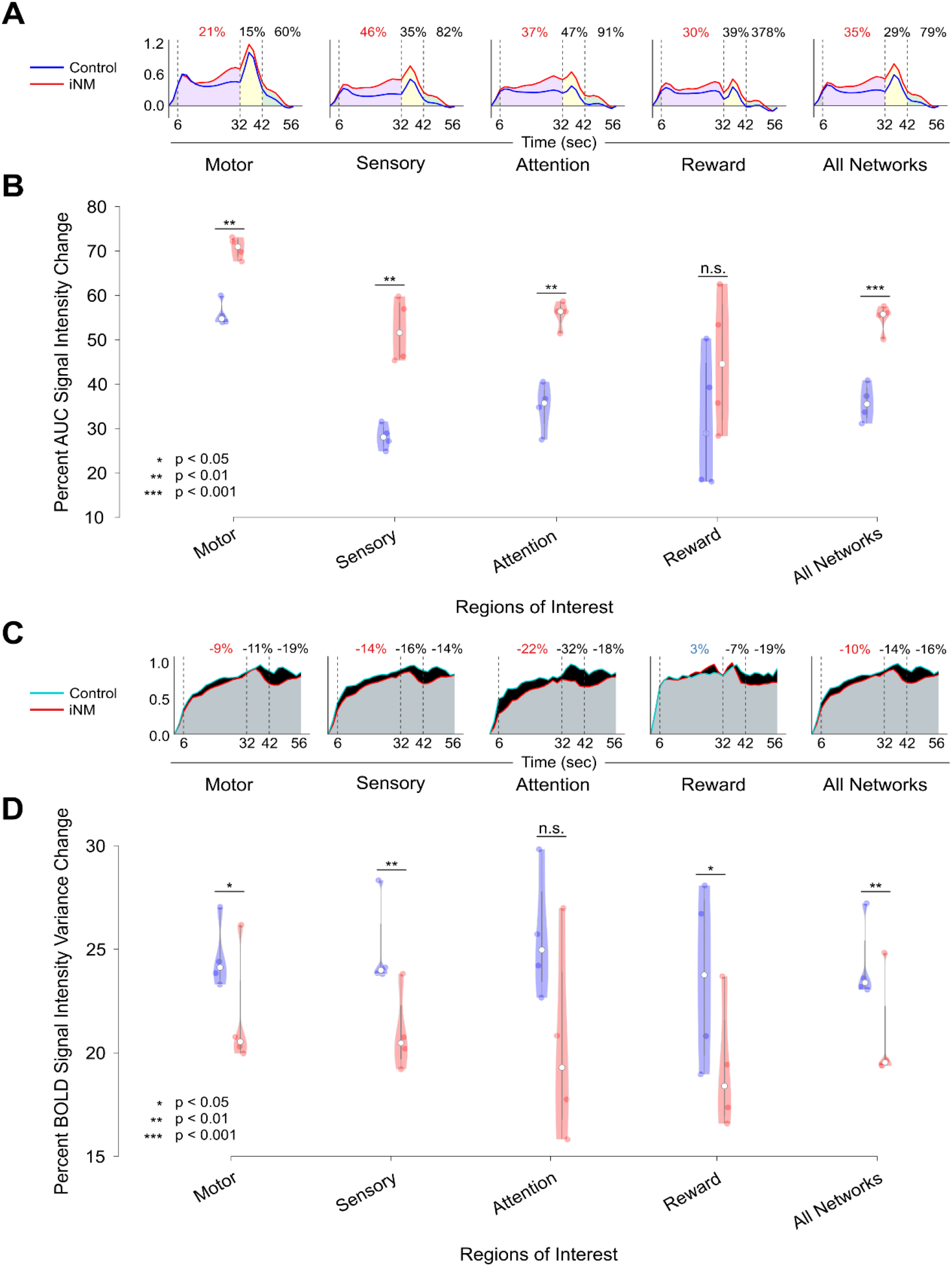
iNM (red) increases mean AUC change in BOLD intensity change and reduces its variance compared to control-no iNM (blue) condition for each network and directions, as a function of time. **A**. The shaded areas (AUC) illustrate cortical direction selectivity during: TMSC (purple, 6 – 32 seconds); swallow (yellow, 32 – 42 seconds); baseline-tongue at rest (green, 42 – 56 seconds). iNM effects transfer to swallow and baseline-tongue at rest blocks during which no iNM is applied. **B**. Significance testing across networks and directions (with dots indicating direction-specific values) showed that iNM differed significantly from control (p < 0.001) in all networks, except in the reward, where no significant differences were observed. **C**. The shaded areas (AUC) signify the same intervals noted as under 4A. Negative AUC values show iNM decreased the BOLD magnitude variance compared to control for all networks, except the reward which showed an increase in the variance. ROIs absent in a specific direction were filled with zeros, results remained unchanged when replaced with random values from a normal distribution (mean = 0, SD = 0.05). iNM effects transfer to swallow and baseline-tongue at rest blocks during which there is no neuromodulation applied. D. Significance testing of variance magnitude across networks and directions (with dots indicating direction-specific values) showed that iNM differed significantly from control (p < 0.001) in all networks except in the attention network, where no significant differences were observe.

When averaged across networks and directions, AUC increased by 35%. Direction-specific changes in AUC and d′ were: 27% and 60.8% (left), 35% and 62% (right), 38% and 63% (up), and 39% and 58.5% (down), respectively (**Figure SI-8A**). During swallow and baseline tongue-at-rest periods—when iNM was not delivered—AUC increases were still observed across directions and networks (except for the reward network in the left direction), suggesting carry-over effects extend their impact beyond active iNM (**Figures 4, SI-8A, SI-8B**). These findings demonstrate that iNM consistently enhanced the magnitude of neural signals within the TMSC network. Increases in the variance of the magnitude were observed in the right tongue direction for the motor (7%), sensory (1%), and reward (12%) networks, with only a 1% overall increase across all ROIs and networks.

Across each and all directions and networks during the swallow period when no iNM was provided, there was a 9% to 18% decrease in the AUC BOLD intensity variance compared to the control. Similarly, during the baseline-tongue at rest period, there was also an 11% to 22% decrease in BOLD intensity variance, for each and all directions and networks. Thus, INM can have a carry-over effect in enhancing TMSC, even when no iNM is applied **(Figures 4, SI-8A—8B)**.

#### Temporal hierarchy shows networks’ engagement as shown by peaks for each region

To examine the temporal hierarchy of activation, we sorted the peak times of each region and identified the following sequence from earliest to latest (**Figure 5C**): 1. the right amygdala (~7 seconds); 2. attention-related regions, including the right medial and middle frontal gyri (8.2 and 8.4 seconds, respectively); 3. several sensory regions, including bilateral S1 (8.6 and 12 seconds), interleaved with the cingulate cortex (10.5 seconds) and bilateral inferior temporal lobules (12.7 and 14 seconds); 4. bilateral anterior cerebellar motor regions (14 and 14.7 seconds); 5. additional sensory areas such as the bilateral thalamus, right inferior parietal lobule, and left insula and claustrum (15.6–15.8 seconds); 6. motor-related regions including bilateral basal ganglia, bilateral inferior frontal gyri, and bilateral primary motor cortices (16–29 seconds); and 7. later-peaking sensory regions such as the right cingulate, right insula and claustrum, and left inferior parietal lobule (16–29 seconds). The hierarchy of peak times for each network’s region within each TMSC direction are shown in **Figure 18**. This temporal ordering of regions by their earliest peak time — from limbic and attentional areas to sensory integration zones, followed by motor execution regions, with a culmination in late sensory re-engagement — highlights the hierarchical and cyclical sensory recruitment of effective TMSC.

**Figure 5.**
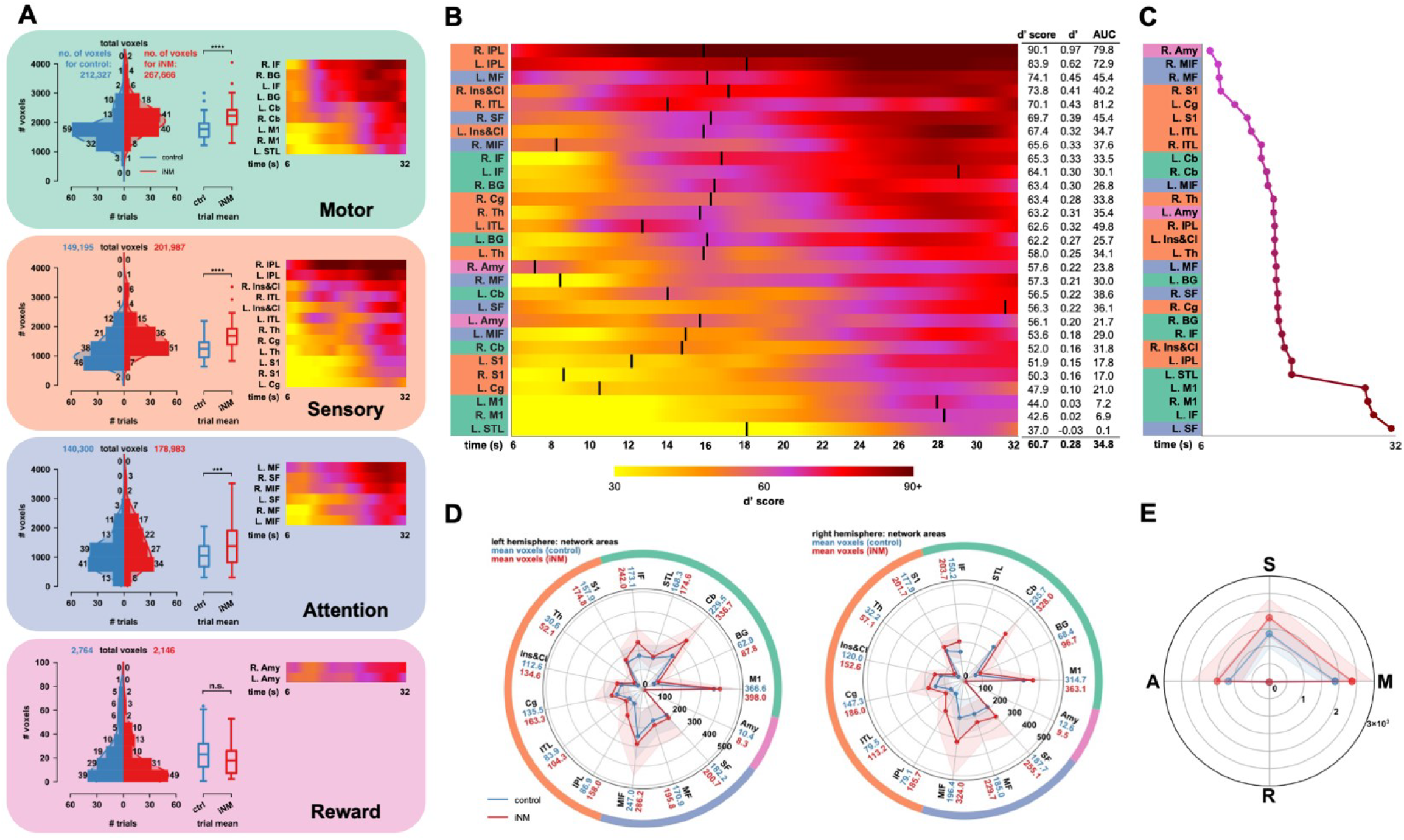
iNM increases the BOLD signal in both magnitude and spatial extent. **(A**) (*left*) For all ROIs within each network (motor, sensory, attention, or reward) under control (blue) and iNM (red), the total number of voxels compromising the subset of ROIs for each trial (subject and direction) is shown in the histogram and kernel density estimate, and summarized in the adjacent box plots (*right*). The means of the d’-generated scores and the percent BOLD signal AUC intensity change are presented in the heatmaps for each region, categorized by network. Red (iNM) and blue (control) indicate the mean d′ score of the BOLD response for each time repetition, averaged across all trials, comparing the iNM and control conditions. (**B**) The same ROIs as in panel A are listed in descending order of overall *d*′ score, across networks and time. Each ROI is color-coded to indicate its corresponding network, as shown in A. The table displays mean d’ score, d’ value, and AUC for each region during the tongue movement interval (6-32 seconds). (C) *d*′ score peak times of each ROI. (D) & (E) Under iNM, the number of voxels within each ROI in the motor (M), sensory (S), attention (A), and reward (R) networks increased relative to the control condition. The solid lines and shaded regions indicate the mean and standard deviation for the control (blue) and iNM (red) conditions, respectively.

#### Consistent Strengthening Within and Across Motor, Sensory, and Attention Network Interactions by iNM Across Tongue Motor Directions

To quantify how iNM changes the stability of latent brain states (e.g., TMSC-dominant vs. swallow-interference states), we employed an unsupervised, autoregressive hidden Markov Model (ARHMM),^48^ which can infer the latent brain states and directional information flow from BOLD signals during TMSC. The algorithm identified two dynamic network states **(Figures 6A, 6B, 6D)**: 1. Dominant state associated with TMSC **(Figures 6A, 6B, 6C)**; and 2. Non-Dominant state associated with swallowing. In the iNM condition, the dominant brain state was upregulated by 15.5% of trials in the TMSC block (97% of trials iNM vs 82% of control trials); 66% of trials in the swallow block (83% vs 50%) and by ~21% of trials in the baseline-tongue-at-rest block (97% vs 80%). ARHMM shows that iNM silenced the non-dominant state from 18% of trials under control down to 3%; i.e., ***an 83*.*3% reduction in swallowing-related interference during the TMSC interval under iNM***. Kernel density estimation (KDE) applied to binned data from each iNM and control block, generated a smoothed, non-parametric estimate of the probability distribution, revealing that 646 trials under iNM and 356 under control were classified as the TMSC-dominant state. Swallow-non-dominance interference during TMSC occurred in only 16 iNM trials, compared to 143 under control, highlighting iNM’s stabilizing effect on TMSC brain states.

**Figure 6.**
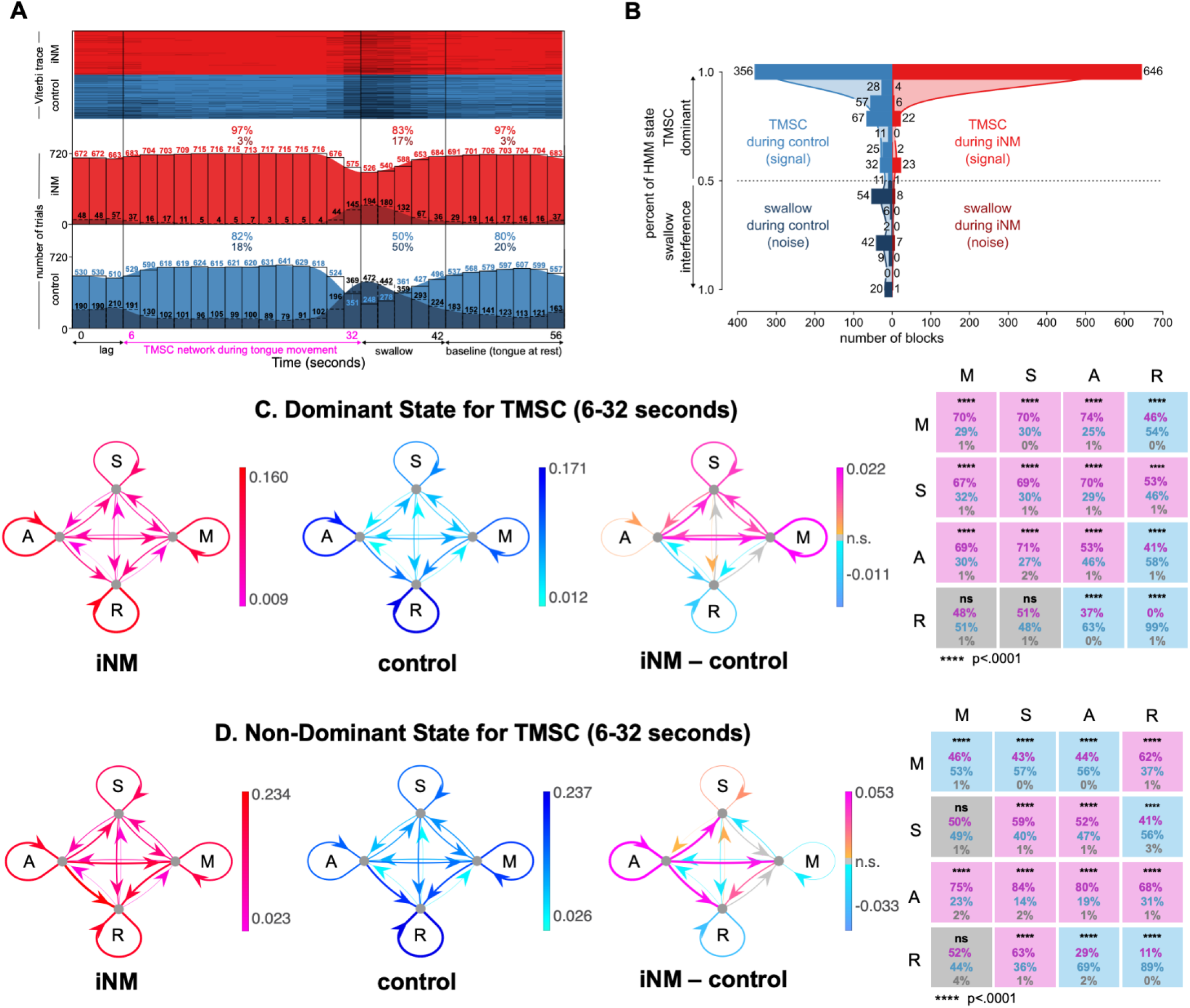
Effect of iNM on network interactions across all tongue motor control directions elucidated via dynamic causal modeling. **A**. We selected the timeseries from the brain areas that select for tongue movement. Our autoregressive hidden Markov model (aHMM) was fit to the brain areas that selected for tongue movement (6-32 seconds). Model inference was then performed on the entire timeseries (0-56 seconds). (top) Viterbi trace for iNM (red) and control (blue) trials that characterize each brain state during TM, swallow, and baseline intervals for each condition. (bottom) Histogram of the number of trials corresponding to TMSC (light red/blue) or swallow (dark red/blue) brain state for each time repetition, divided into the iNM (red) and control (blue) conditions. Percentages correspond to the fraction of trials within each brain state across the TM, swallow, or baseline intervals. **B**. The histogram compares the signal (TMSC state, red/blue) and noise (swallow state, dark red/dark blue) by displaying the number of trials in each bin that correspond to each of the two states (total trials (iNM vs. control): 727 vs. 525 (TMSC state), 15 vs. 195 (swallow state)). Shaded curves represent the kernel density estimate (KDE) fit to the binned data for the iNM and control conditions. The KDE was calculated by the summation of normal distributions centered at each data point, followed by normalization, to generate a smoothed estimate of the histogram bins. KDE bandwidth was estimated via Scott’s method, and boundary corrections at 0 and 1 employed reflection of the data across the boundaries. **C-D**. For the dominant (TMSC) or non-dominant state, the strength of ROI-to-ROI connections within networks (M = Motor network, S = Sensory network, A = Attention network, R = Reward network) is quantified as the mean absolute value of the autoregressive coefficients during the iNM (fuchsia/red) or control (light/dark blue) condition, or the difference in absolute values between conditions (orange/fuchsia and light/dark blue indicate increased/decreased during iNM, respectively). At far right, significant increases (fuchsia), decreases (light blue), or insignificant differences (gray) in these network-to-network interactions are shown for iNM compared to control. Percentages indicate the fraction of region-to-region interactions within each network-to-network connection that are increased (fuchsia), decreased (light blue), or insignificant (gray) in iNM compared to control. Correction for multiple comparisons used an FDR of 0.05.

ARHMM also estimated functional connectivity between ROIs within each network for both dominant and non-dominant states. Significance testing revealed ROI connections that were either upregulated or downregulated under iNM and control conditions (**Figures 6C, 6D**). During the dominant state, iNM significantly increased connectivity within and across motor, sensory, and attention networks, ranging from 53% to 74% across all TMSC-selective regions (p<0.0001; **Figure 6C**), while for the reward network, there dominant state decreased under iNM compared to control. The ARHMM also identified consistent patterns in the prevalence of dominant versus non-dominant states across TMSC movement directions: the dominant state was present in 96% of iNM trials versus 88% under control for the left direction (**Figure SI-9**); 96% vs. 79% for the right (**Figure SI-10**); 95% vs. 78% for the up (**Figure SI-11**); and 97% vs. 81% for the down direction (**Figure SI-12**). During the non-dominant state, the ARHMM revealed a statistically significant reduction in the proportion of trials under iNM compared to control (p<0.0001; **Figures 6A, 6B**). Specifically, for the left direction only 4% of trials were characterized as non-dominant under iNM versus 12% under control (**Figure SI-9**); for the right direction, 4% under iNM versus 21% under control **(Figure SI-10)**; for the up direction, 5% under iNM versus 22% under control **(Figure SI-11)**; and for the down direction 3% under iNM versus 19% under control (**Figure SI-12**). Across all movement directions, TMSC dominance ranged from 95–97% of iNM trials, accompanied by reduced prevalence of the competing, swallow-associated state that ranged from 3-5%. Overall, iNM reinforced a new network -- anchored in motor, sensory, and attention areas -- in its magnitude, spatial extents, and stability that controlled tongue movement and direction.

## Discussion

### Role of individualization in targeting tongue representations

Biological systems are inherently complex, in part due to their nonlinearity, which gives rise to the variability that defines their function.^49^ ***This inherent variability necessitates modeling each system individually***, that can precisely capture subject-specific neural dynamics. The iNM intervention constructs subject-specific functional and anatomical profiles from fMRI data (**Figure 2**). This process generates a data-driven representation of an individual’s neural network architecture— a “neurofunctional twin”—that serves as a functional analog^50^ -- across the iNM scans, crucial for the regulation of each individual’s unique TMSC representation. We propose that iNM engages distinct mechanisms depending on lesion location. For suprabulbar lesions, as in neurodegenerative disease, iNM may strengthen intact circuits to bypass damaged pathways and partially restore lost functions. In contrast, for infrabulbar lesions, such as those found in head and neck cancer, iNM is hypothesized to enhance compensatory afferent and efferent signaling by recruiting collateral branches near cranial nerves IX and XII. Second, iNM operates through closed-loop adaptation, using iterative modulation within the fMRI environment to update the participant’s engagement strategy based on real-time neural responses. This mirrors the adaptive updating process inherent in digital twin systems. Third, iNM is designed for prediction and optimization—the individualized functional maps are used to assess which areas engage the most, much like how digital twins refine interventions in engineered systems. Finally, iNM operates as a biophysically grounded intervention: it relies on the MRI’s magnetic field and BOLD signal to measure oxygenation-dependent, quantifiable parameters that directly guide modulation of brain activity. The scanner’s magnetic field interacts with the individual’s vascular and metabolic landscape, dynamically altering local magnetic susceptibility.^51^ iNM guides these subject-specific anatomical and functional high-performing neural circuits by reinforcing and refining them, thus promoting greater selectivity for TMSC precision, bilateral recruitment, and stability.

### iNM enhances the consistency, specificity, and stability of neural activation across tongue motor and sensory control (TMSC) networks

Compared to the control condition, iNM increased BOLD signal magnitude, reduced variance, and expanded spatial representation across motor, sensory, and attention networks. Direction-specific activation patterns became more distinct under iNM, as revealed by SVM classification, which reached 94.3% accuracy for tongue movement versus baseline (**Figure 3, SI-1 - 3**). These effects reflect enhanced signal-to-noise ratio (SNR), improved functional specificity, and more consistent recruitment of individualized TMSC networks. Notably, iNM selectively enhanced relevant circuits while limiting expansion in the reward network, aligning with its targeted motor-sensory focus.

Temporal analyses revealed that iNM effects persisted beyond the stimulation window. Area under the curve (AUC) metrics showed sustained BOLD intensity across functional networks and movement directions, extending into rest and swallow periods (**Figures 4, 6**). Spatially, network coverage increased dramatically—614% in the sensory network, 230% in motor, and 166% in attention—with the largest regional expansions in the bilateral inferior parietal lobule (135% in the right IPL, 82% in the left IPL) and thalamus (77.3% in the right, 70.3% in the left), underscoring their critical role in sensory gating during tongue movement (**Figures 5B, 5C, 5D**). In contrast, the reward network exhibited a 45% decrease in spatial expansion. An overall 10% reduction in the variance of the BOLD magnitude quantified by AUC, indicates improved signal stability and a higher SNR under iNM (**Figure 4C, 4D, 6B**). Notably, iNM exhibits a carryover effect beyond the neuromodulation window, as evidenced by increased AUC extending into swallow and baseline-tongue-at-rest period, even in the absence of iNM. This pattern suggests that iNM not only boosts immediate neural decoding accuracy, stability and adaptive persistence across TMSC directions and networks.

### Bilateral inferior parietal lobule (IPL) engagement underlies adaptive, circuit-specific modulation during iNM

IPL emerged as a central hub governing iNM-induced sensorimotor adaptation, showing no significant engagement under control conditions. Under iNM, IPL exhibited strong discriminability (right d′ = 90%, AUC = 80%; left d′ = 84%, AUC = 73%) and substantial spatial expansion (right: +132%, left: +82%), reflecting increased sensorimotor demands. Notably, BOLD variance in IPL rose by 53% compared to control, suggesting dynamic prediction-error processing rather than signal instability. This aligns with models of internal state updating and error correction, where increased variability enables real-time recalibration of motor plans. Supporting this interpretation, prior studies link IPL to gamma-band synchronization, forward model correction, and precision-weighted error processing.^52–54^ Temporally, right IPL peaked before left IPL, suggesting sequential sensory integration followed by motor adjustment. These features position IPL as a multimodal integration center crucial for the adaptive modulation of TMSC.

### iNM engages a temporally coordinated network supporting anticipatory control, sensory integration, and motor refinement

Early peak activation in the right amygdala, followed by medial inferior frontal (MIF) regions and bilateral S1, suggests anticipatory control and proprioceptive tuning precede motor execution. Amygdala-MIF interactions may shape task salience and internal state monitoring, while early S1 activation (right: 8.6 s; left: 12 s) likely supports real-time proprioceptive feedback. Bilateral M1 engagement occurred later (~28 s), reinforcing a sensory-to-motor flow. Concurrently, sequential activation of cingulate cortices and inferior temporal lobes indicates layered cognitive control, imagery, and motor planning. Peak engagement of the cerebellum (left: 13 s; right: 14 s) and basal ganglia (left: 16 s; right: 16.4 s) underscores their role in adaptive, time-locked regulation of tongue movement. Together, these findings reveal a distributed, temporally coordinated network recruited under iNM to support anticipatory evaluation, error correction, and precise TMSC control.

### Temporal causal modeling via unsupervised ARHMM

Unsupervised autoregressive hidden Markov modeling (ARHMM) revealed that iNM consistently stabilized the TMSC-dominant brain state, suppressing swallow-related interference, while it enhanced TMSC state persistence not only during tongue movement, but also duirng swallowing, and rest intervals. During TMSC blocks, 97% of iNM trials were classified as TMSC compared to 82% in the control, with similar dominance observed during swallow and baseline periods, indicating sustained TMSC network engagement and carry-over effects. This was accompanied by reduced BOLD signal variance, reflecting improved neural stability and error minimization—hallmarks of efficient task selection and experience-dependent plasticity. Notably, iNM strengthened within- and across-network connectivity in motor, sensory, and attention networks (53–74%), while suppressing reward-network interactions, consistent with its targeted rehabilitative focus.

These effects suggest that iNM promotes both functional specificity and adaptive persistence by optimizing TMSC stability without disrupting flexibility. This consolidation of task-relevant states across time windows supports a mechanistic foundation for neurorehabilitation, where lasting gains depend on generalization beyond trained tasks.^55–57^ Functionally, iNM acts as a real-time, adaptive brain-computer interface—precisely modulating circuit-level dynamics through individualized, hemodynamically coupled feedback. Its architecture mirrors key features of a digital twin: continuous tracking, personalized control, and dynamic adjustment of brain networks based on real-time input. As a biologically grounded, closed-loop system, iNM reorganizes TMSC circuits in a durable, scalable manner—laying the groundwork for restoring sensorimotor function in neurological disorders marked by diffuse or asymmetric damage. By accessing the full brain and adapting to individual connectivity patterns, iNM offers a pathway toward personalized, high-precision neuromodulation for functional recovery.

### Future Directions: additional iNM applications for motor and sensory control

Beyond its application in radiation-induced cranial neuropathy (RICN), iNM holds promise for other forms of cranial and peripheral neuropathy. CCN can also manifest following anterior cervical discectomy and fusion (ACDF) surgery, where cranial nerve and tongue motor deficits may arise secondary to surgical or inflammatory insult. Although ACDF is a common procedure -- comprises 80% of cervical spinal surgeries -- and can safely treat cervical disc herniation with a 70% success rate,^58,59^ it carries risks, such as esophageal perforation, laryngeal nerve palsy, dyspnea, dysphonia.^60–69^ Dysphagia in ACDF, affects 34–38% of patients with 25.5%, developing chronic symptoms that lead to tongue muscle atrophy.^60,61,70– 77^ impacting quality of life, and which in severe cases can be fatal.^78^ Similarly to CCN in head and neck cancer survivors, effective treatments remain limited,^27,79–88^ iNM may offer a promising neurorehabilitation strategy following ACDF surgery.

Beyond CCN, iNM can be used for peripheral neuropathies experienced after breast cancer (BCA) surgery or radiation, using the same principle of strengthening motor and sensory networks^89^. Despite high survival rates, 20–60% of BCA patients develop post-mastectomy pain syndrome (PMPS), a debilitating neuropathy following lymph node dissection affecting the thorax, axilla, and upper limb,^90–92,95–97^ as a result of adhesions or hematomas involving the intercostobrachial nerve.^98–100^ This pain interferes with inspiration, expiration, and trunk movement, accompanied by severe depression, as 54% of affected survivors report disruptions to employment and access to healthcare.^98–107^ Similarly to CCN, the need for a safe, noninvasive treatment is apparent here, as well.^108–111^ iNM could be adapted to target sensorimotor circuits involved in arm strength, proprioception, and pain modulation^89^. The BCA survivorship space underscores the potential of iNM to move beyond orofacial applications, enabling targeted modulation of sensorimotor circuits involved in upper limb strength, movement, proprioception, and pain regulation.

Together, these applications illustrate the broader potential of iNM to transform neurorehabilitation. iNM enables precise, circuit-specific modulation by capitalizing on the coupling between hemodynamics – which reflects neural dynamics -- and the MRI’s field. By integrating adaptive control, real-time neuroimaging, and millimeter-precise targeting, iNM serves as a neurofunctional model -- individualized, and predictive -- that updates the brain’s TMSC networks, rather than a fixed therapeutic tool. As such, iNM can serve as a scalable platform for restoring motor and sensory function across a wide range of neurologic, post-surgical, and post-radiation conditions through precision-guided intervention: i.e., *it can access each subject’s entire brain – a key factor to successfully manage unique symptoms to each patient, based on the afferent or efferent nerves lesioned, and thus, brain networks involved*. In doing so, iNM not only advances the field of brain-computer interfacing and neurofunctional twin-inspired modeling, but also lays the foundation for a new generation of non-invasive, individualized, neuromodulation therapies.

## Acknowledgements

This study is funded by grants to T.D. Papageorgiou: McNair Medical Institute; Robert and Janice McNair Foundation; Mission Connect, IP Award - The Institute of Rehabilitation Research; Center for Alzheimer’s and Neurodegenerative Disease; Jan and Dan Duncan Neurological Research Institute; Naman Basic Science Faculty Award; Fight for Sight; NIH-T32, NEI; NIH-T32, NINDS, Breast SPORE – Development Research Project Award.

TDP wishes to thank Dr. Sameer A. Sheth, Professor – Cullen Foundation Endowed Chair of Neurosurgery and Vice Chair of Research Neurosurgery at Baylor College of Medicine for his support of this study.

TDP gratefully acknowledges the legal guidance provided by the intellectual property team Kevin Keeling and James P. Cargas, whose expertise led to the successful patenting of the iNM platform (Patent No. 19 753 851.5, WIPO Publication No. WO 2019/160754).

