## Supplemental Figures and Tables for "Individualized AI-driven neuromodulation enhances tongue motor and sensory control networks: a precision intervention for neurorehabilitation in cancer survivors and patients with neurodegenerative disease"

**Author Contributions:** T.D.P. developed iNM, conceptualized, designed research; T.D.P, D.H. performed study experiments; T.D.P., J.W., A.A., S.R., D.H., R.H., and E.F. contributed analytic tools; T.D.P.  J.W., A.A., S.R., D.H., R.H. analyzed data; T.D.P., EF, J.W., A.A., K.H., E.M.R., E.M.S., AMT, S.G.H., R.H., C.N. interpreted findings; T.D.P. and E.F. wrote the paper.

**Competing Interest Declaration --** Patent granted to Dr. T. Dorina Papageorgiou & Dr. E. Froudarakis: No. 19 753 851.5, WIPO Publication No. WO 2019/160754

**Abstract (word count: 250 – limit: 250)**
Precise modulation of brain networks responsible for tongue motor and sensory control (TMSC) is critical for restoring functions, such as speech and swallowing in neurodegenerative disease or in treatment-induced chronic cranial neuropathy. We present an individualized, AI-driven fMRI neuromodulation (iNM) platform that adaptively targets subject-specific TMSC networks in real time. To enhance iNM precision and encodability —critical for neurorehabilitation—we mapped each healthy participant's individualized TMSC selectivity network, creating a subject-specific TMSC digital twin. iNM increased signal strength, spatial expansion, and consistency across motor, sensory, and attention regions, while it reduced signal variability. The bilateral inferior parietal lobule emerged as key sensorimotor integration hub, as it exhibited exclusive activation under iNM along with highest discriminability, and largest spatial expansion. iNM also significantly strengthened and expanded motor, sensory, and attention-related networks — medial-middle frontal areas, insula-claustrum, S1, M1, basal ganglia, motor cerebellum, and inferior temporal— supporting interoceptive and proprioceptive-motor integration. Machine learning and unsupervised hidden Markov modeling revealed that iNM enhanced the decodability and stability of TMSC-neural states, while it suppressed competing swallow-neural state interference. Notably, the iNM effects extended beyond the neuromodulation window, indicating functional persistence—a key requirement for rehabilitation. iNM reconfigured TMSC networks by strengthening cortico-subcortical connectivity and adaptive circuit dynamics. Our findings show iNM as a non-invasive, personalized intervention capable of selectively enhancing sensorimotor control with high spatiotemporal specificity. By demonstrating mechanistic network-precision and functional carryover, iNM offers a promising intervention for individuals with limited treatment options, including head and neck cancer survivors and early-stage neurodegenerative disease patients.

### Extended Data Figures

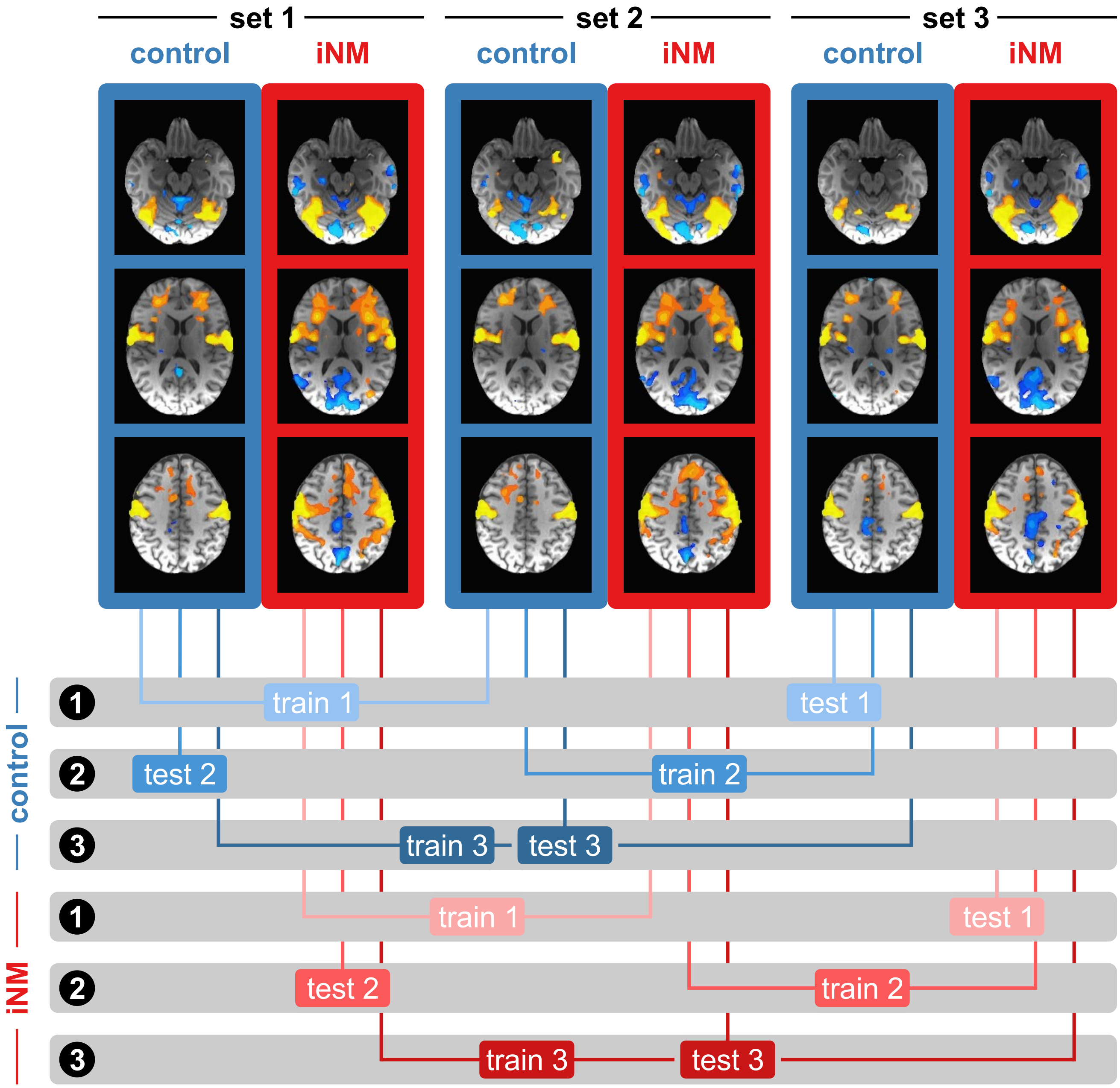


**Extended Data Figure 1. Training and testing SVM permutations using a 3-fold cross-validation design.** SVM classification accuracy for each permutation was generated by computing the mean accuracies from the three corresponding test datasets.

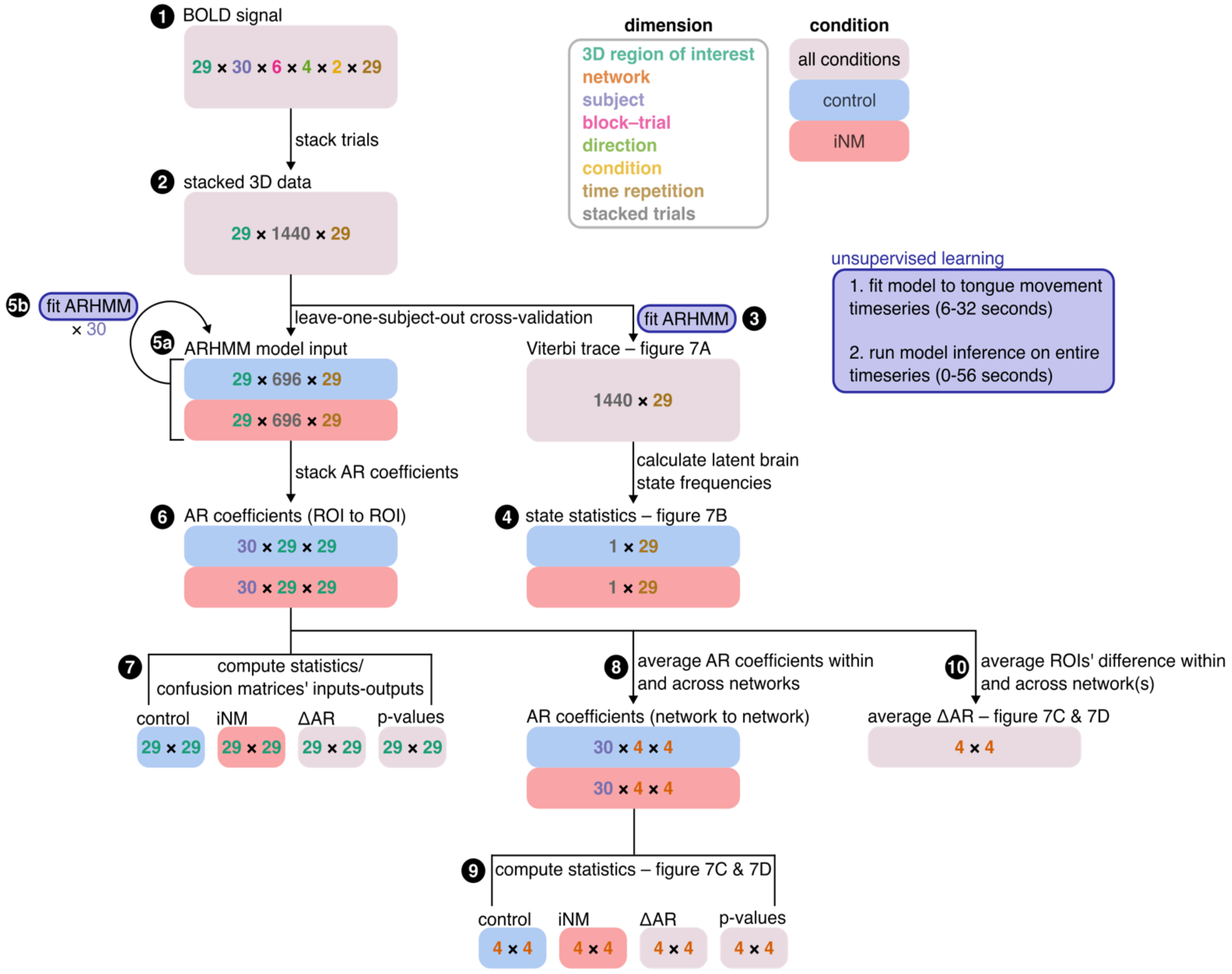


**Extended Data Figure 2: Step-by-step overview of autoregressive hidden Markov model (ARHMM) analysis.** BOLD signal data (**1**) was organized by stacking trials into a single dimension (**2**) on which ARHMM analyses could be performed. The ARHMM was fit only to the timeseries corresponding to tongue movement, but model inference, via the Viterbi trace, was performed on the entire timeseries (**3** and **5**; see inset in purple box). First, the ARHMM was fit to both conditions simultaneously (**3**), and frequencies of the latent states were then calculated for trials corresponding to control and iNM conditions separately (**4**). Next, leave-one-subject-out cross-validation was performed on the stacked dataset. All trials from one subject were held out (**5a**), while ARHMM models were fit to data from the remaining twenty-nine subjects under each condition (**5b**), resulting in a total of thirty ARHMM models for both the control and iNM conditions. From the autoregressive (AR) coefficients for these models (**6**), the average confusion matrices representing functional connectivity between regions-of-interest (ROIs) were calculated (**7**). AR coefficients were subsequently grouped by network (**8**; i.e. motor, sensory, attention, reward), and confusion matrices representing network-to-network interactions were similarly computed (**9**). Lastly, within each network, the sum of differences between the coefficients corresponding to control and iNM conditions were calculated (**10**).

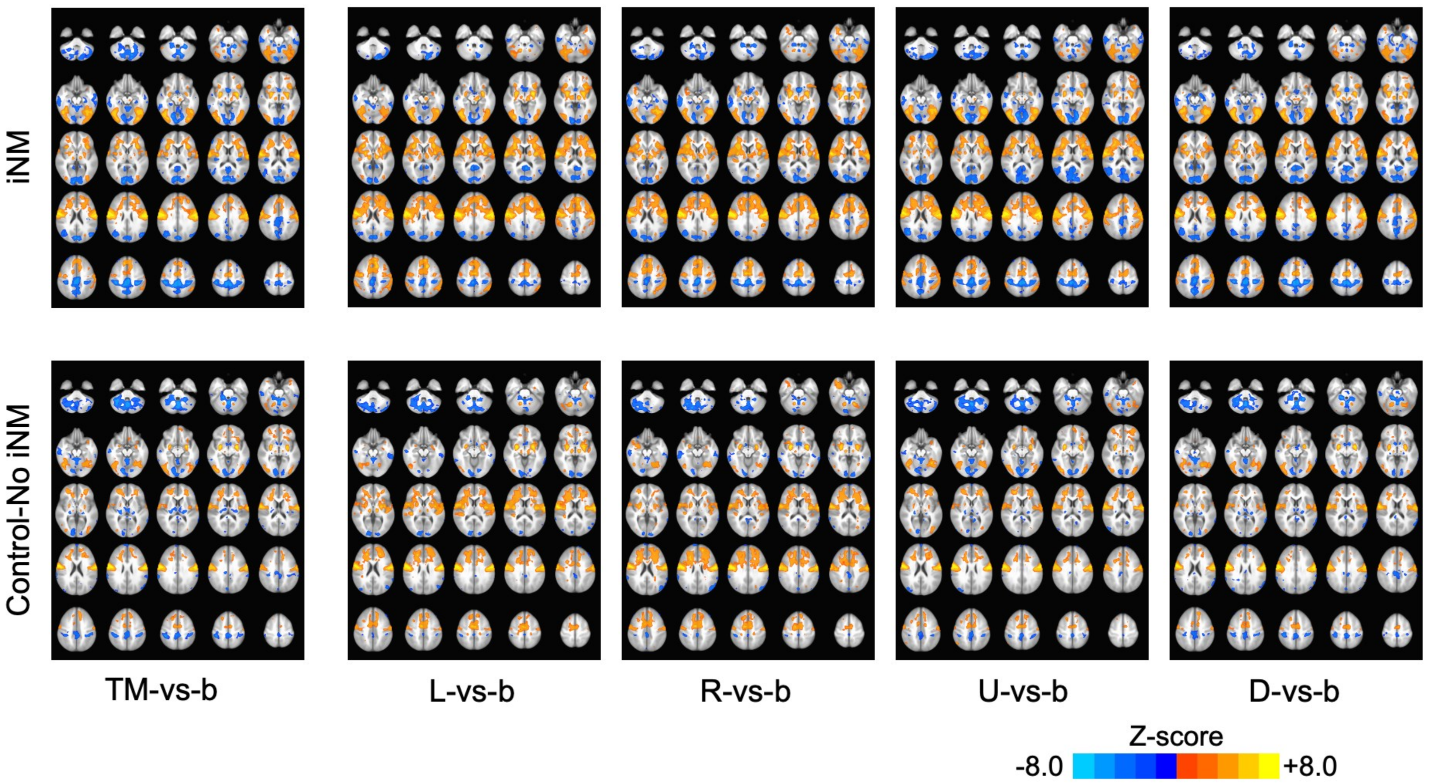


**Extended Data Figure 3. Cortical maps show that iNM increases consistency in the signal intensity and spatial extent compared to the control-no iNM condition using linear machine learning -- support vector machine analysis.** SVM-generated activation maps for each tongue motor control direction are shown for the iNM condition (top) and the control (no iNM) condition (bottom). Maps are thresholded at a voxelwise p-value of 0.001 (FDR-corrected, $q \leq 0.01$), with a minimum cluster size of >10 voxels

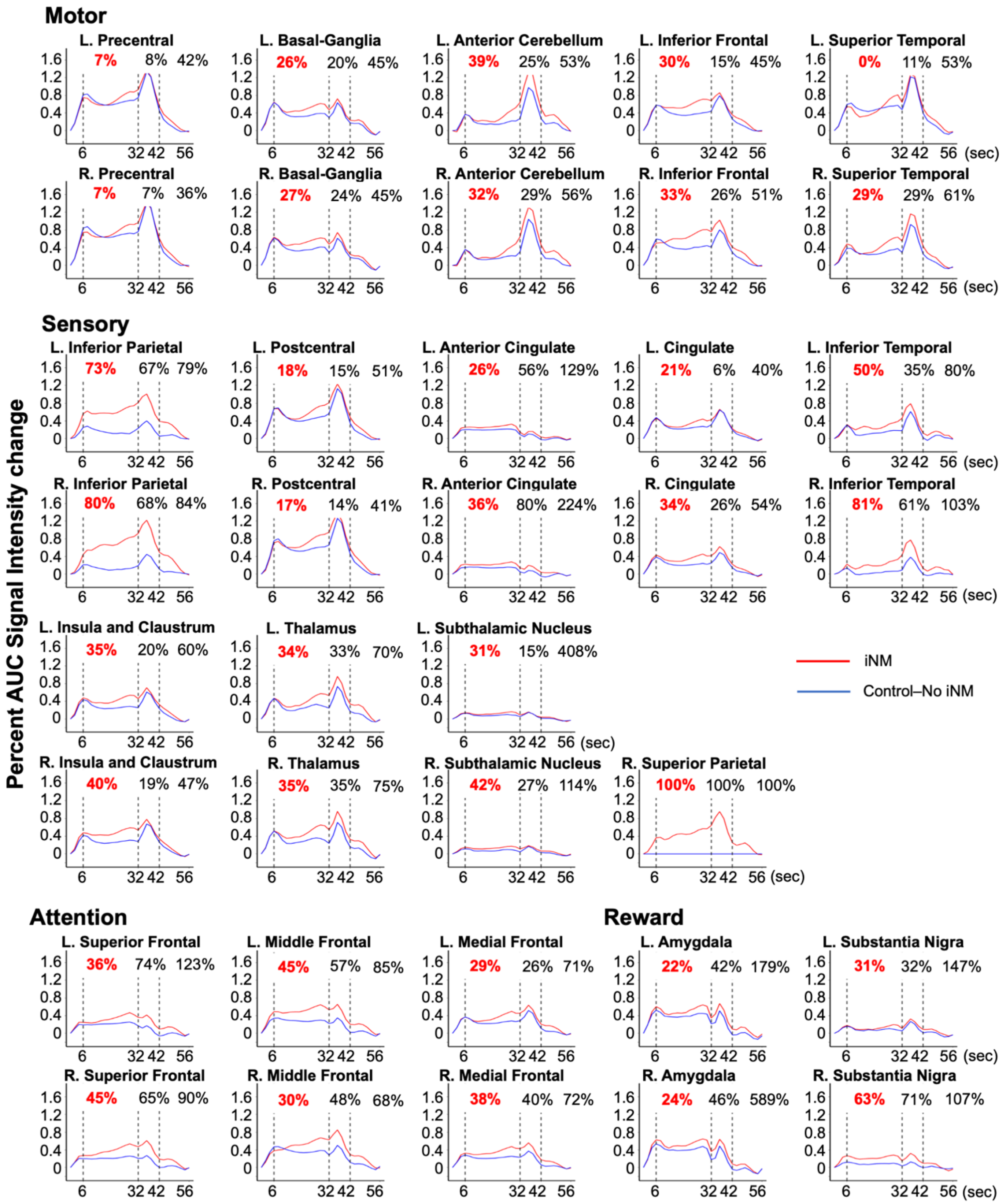


**Extended Data Figure 4. iNM (red) increases mean percent AUC signal intensity change compared to control-no iNM (blue) condition for each ROI within each network**. The mean percent AUC signal intensity change, shown for ROIs categorized by network, illustrate cortical direction selectivity during tongue movement (6 – 32 seconds); cortical selectivity during swallow (32 – 42 seconds); and cortical selectivity during baseline-tongue at rest (42 – 56 seconds). iNM led to increases in AUC ranging from 7–39% in motor, 17–100% in sensory, 29–45% in attention, and 22–63% in reward-related regions, reflecting consistent enhancement across the constituent areas of each functional network.iNM effects transfer to swallow and baseline-tongue at rest blocks during which there is no neuromodulation applied.

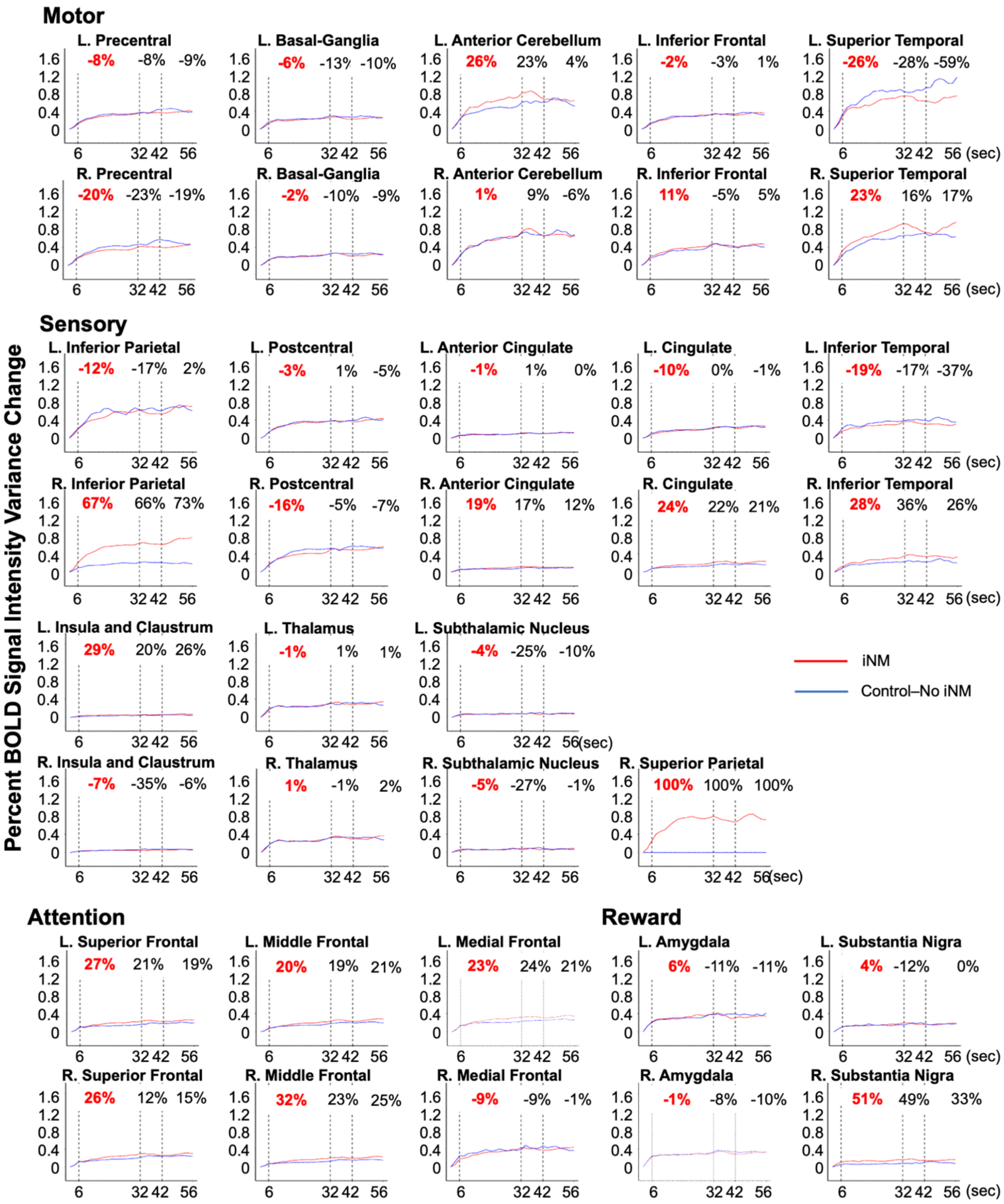


**Extended Data Figure 4b. iNM (red) increases mean percent AUC signal change of the variance magnitude compared to control-no iNM (blue) condition for each ROI within each network.** The mean percent AUC signal change of the variance's magnitude, shown for ROIs categorized by network, illustrate cortical direction selectivity during tongue movement (6 – 32 seconds); cortical selectivity during swallow (32 – 42 seconds); and cortical selectivity during baseline-tongue at rest (42 – 56 seconds). iNM led to increases in AUC ranging from 7–% in motor, 17–100% in sensory, 29–45% in attention, and 22–63% in reward-related regions, reflecting consistent enhancement across the constituent areas of each functional network.iNM effects transfer to swallow and baseline-tongue at rest blocks during which there is no neuromodulation applied.

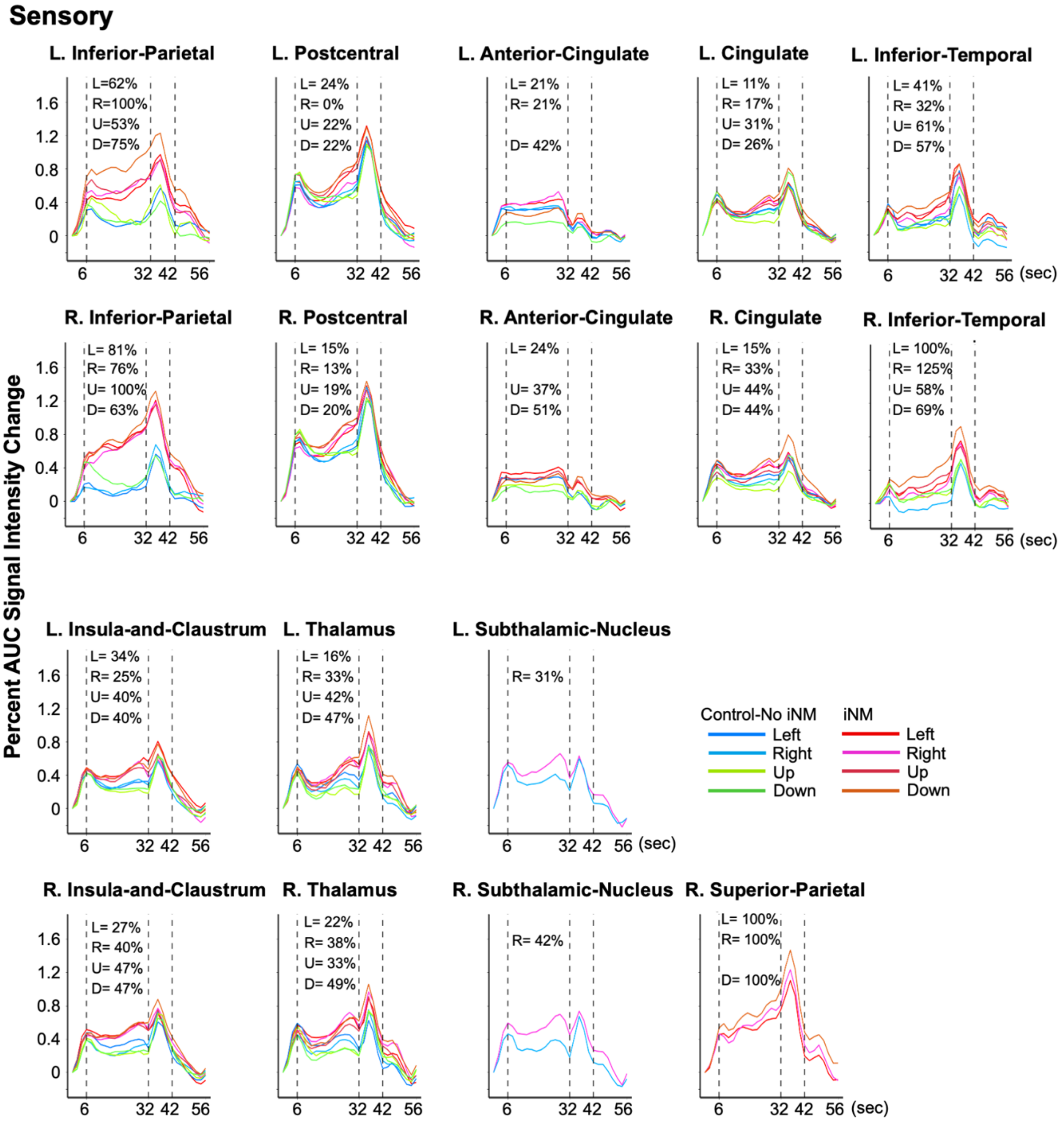


**Extended Data Figure 5a. iNM increases mean percent AUC signal intensity change compared to control-no iNM condition for each cortical direction selectivity during tongue motor control (6 – 32 seconds) within each ROI that comprise the sensory network.**

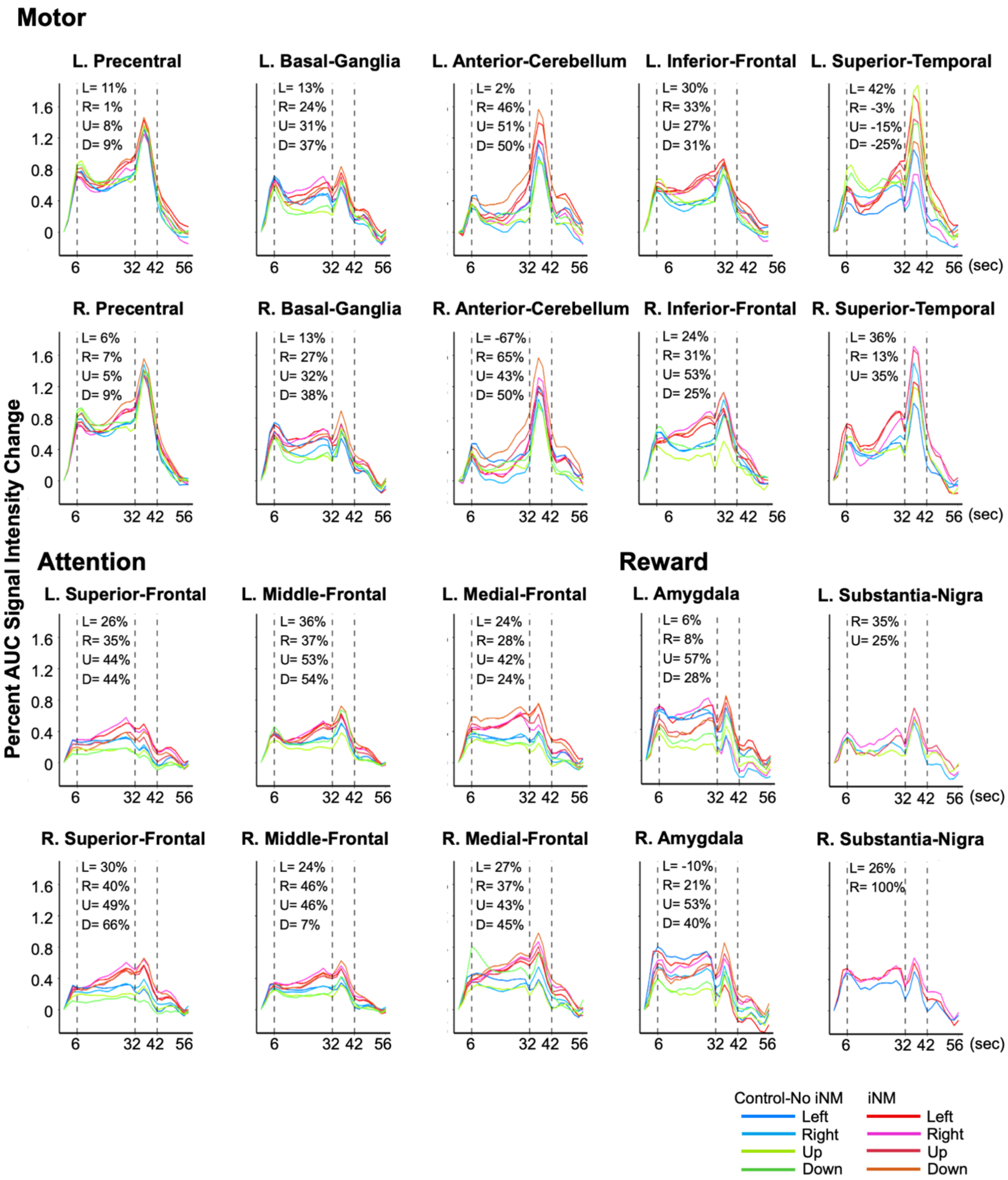


**Extended Data Figure 5b. iNM increases mean percent AUC signal intensity change compared to control-no iNM condition for each cortical direction selectivity during tongue motor control (6 – 32 seconds) within each ROI that comprise the motor, attention, and reward networks.**

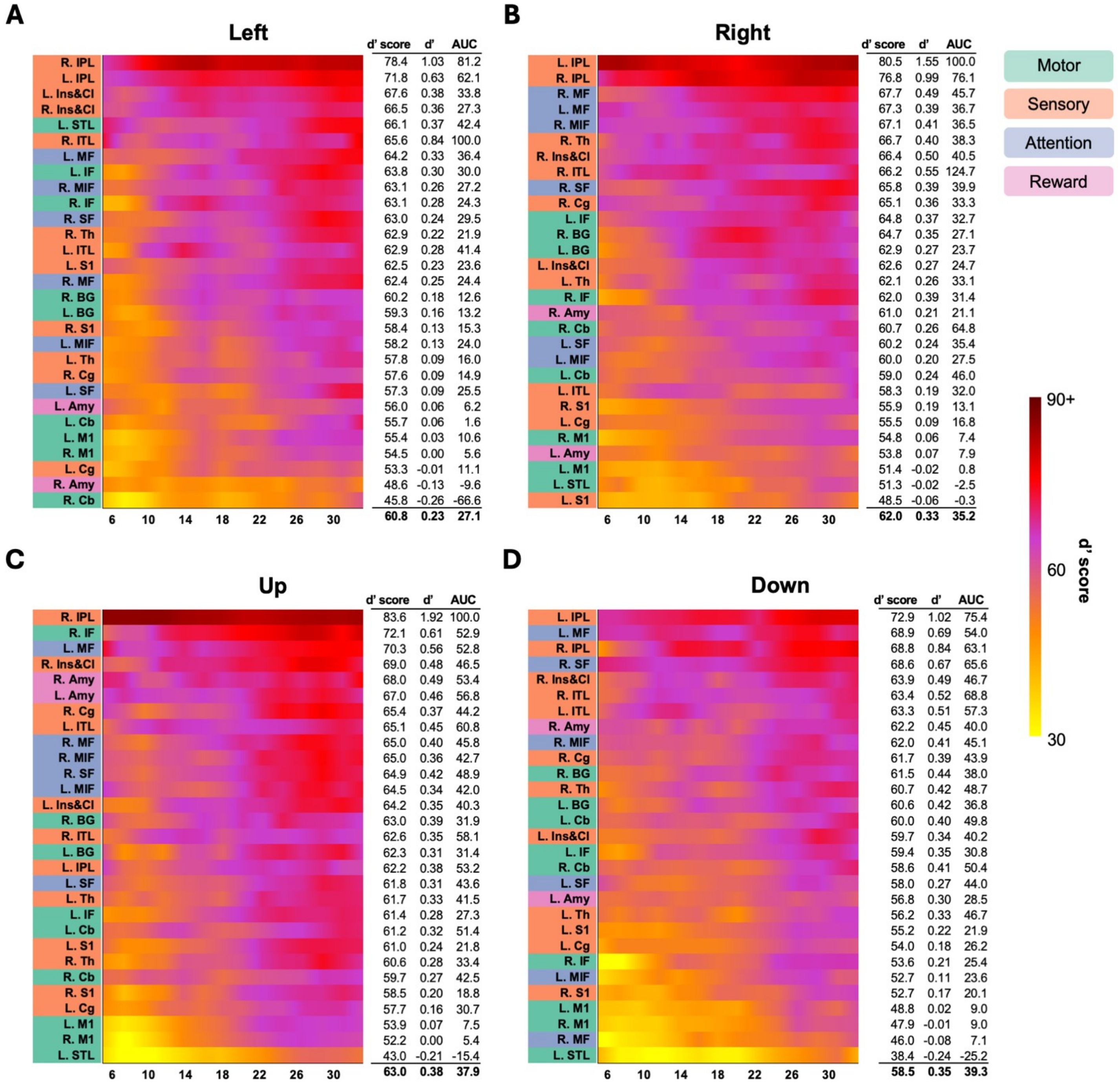


**Extended Data Figure 6: iNM increases the sensitivity index (d'), d' score, and AUC spatiotemporally for TMSC direction selectivity as a function of direction compared to control.** Similar to Figure 5, the heatmaps for the left left (**A**), right (**B**), up (**C**), and down (**D**) directions visually the mean d' score of each region that compares BOLD response between the iNM and control conditions, for a given time repetition. Heatmap figures display the mean d' score, d' value, and AUC during the tongue movement interval (6-32 seconds). Region labels are highlighted according to network.

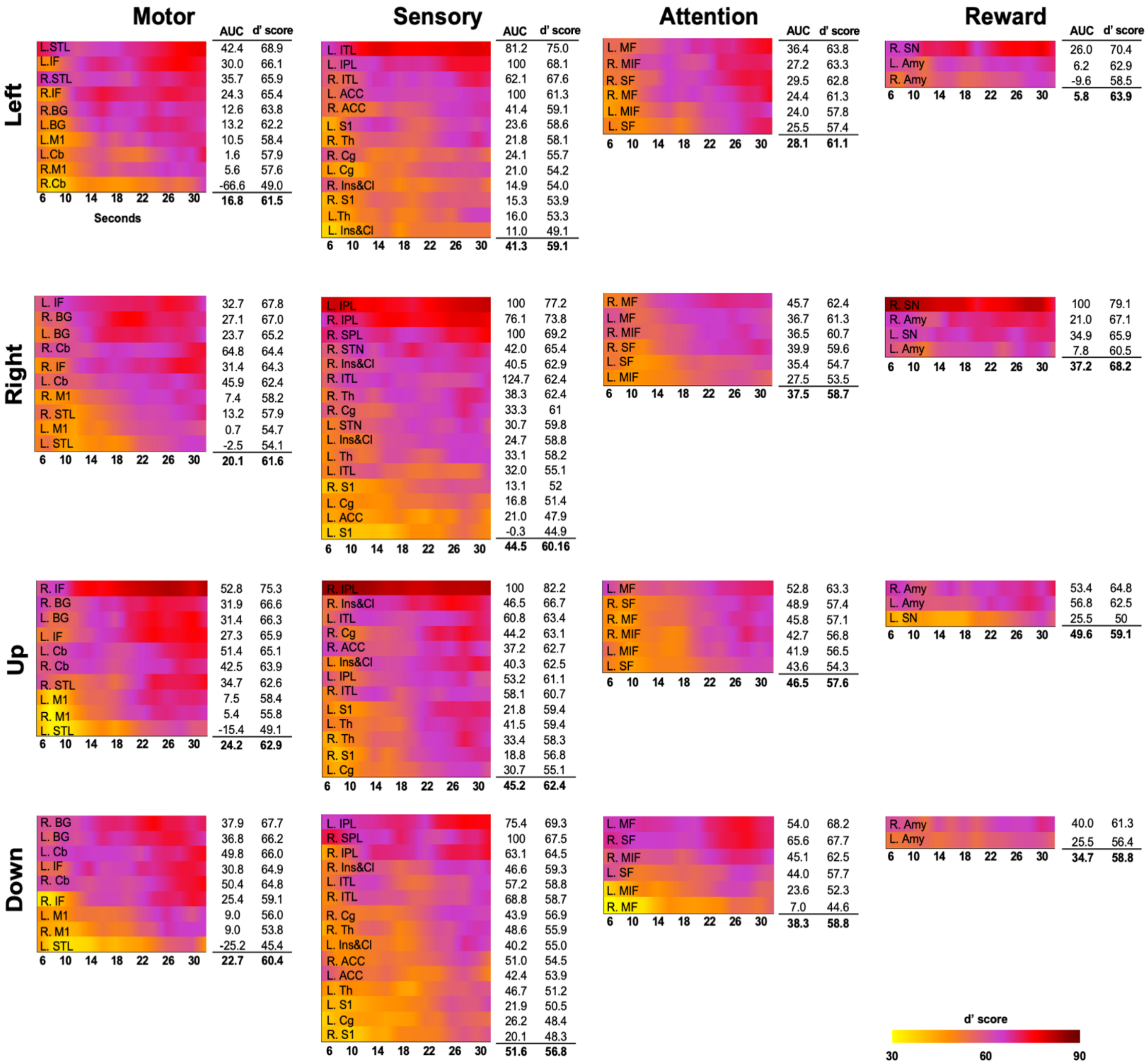


**Extended Data Figure 7: iNM increases the sensitivity index (d') and AUC spatiotemporally for TMSC direction selectivity as a function of network and direction.** Heatmaps of the d' scores over time, as well as mean AUC and d' score per region, are shown as in Figure 6, except here organized by tongue movement direction and network.

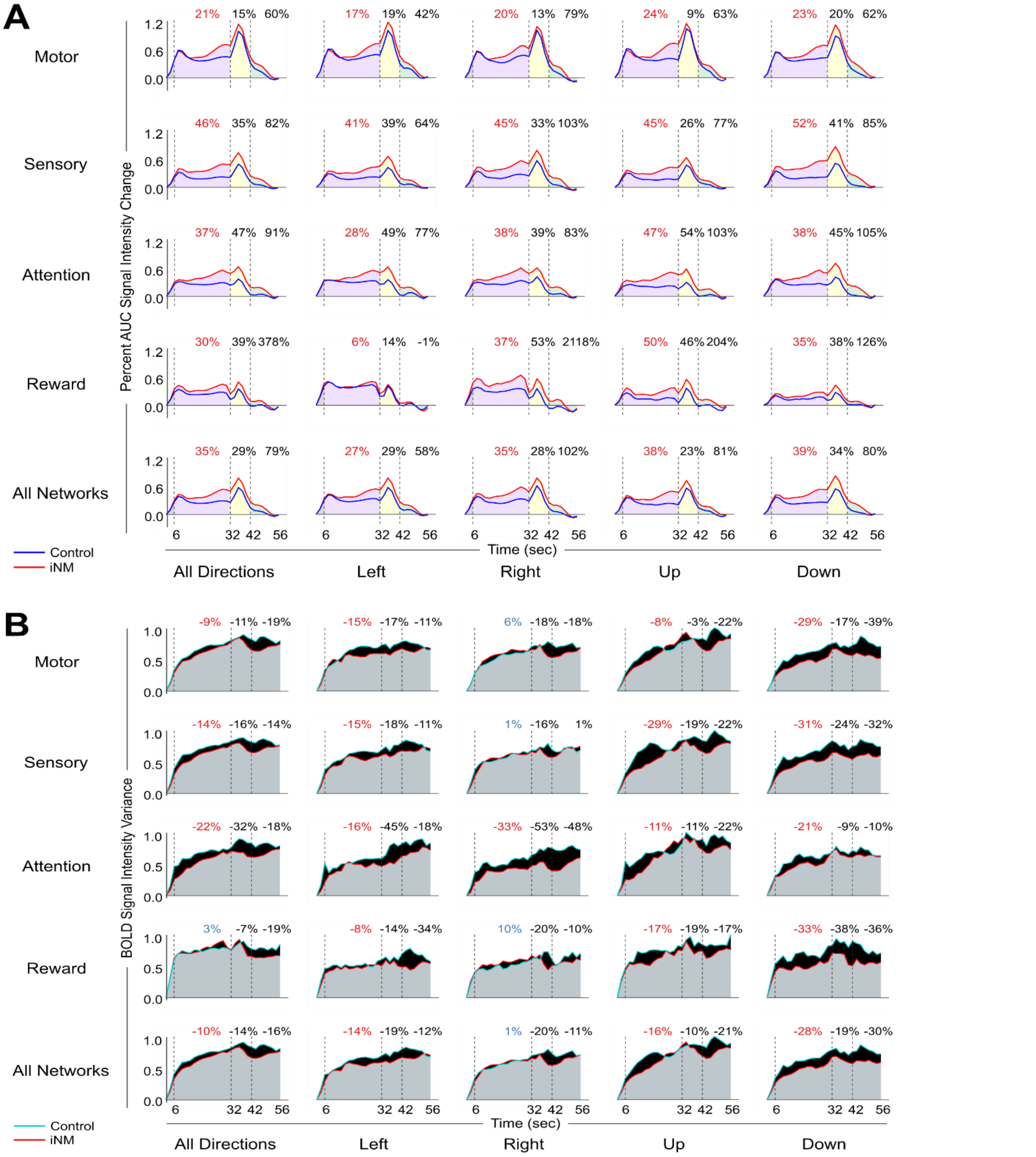


**Figure 8. iNM (red) overall decreases the mean percent AUC change of the BOLD intensity variance compared to the control (blue) condition for most tongue motor control directions and networks, as a function of time**: 1. cortical direction selectivity during tongue movement (6 – 32 seconds) 2. cortical selectivity during swallow (32 – 42 seconds) 3. cortical selectivity during baseline-tongue at rest (42 – 56 seconds). Negative (positive) values correspond to iNM-induced decreases (increases) of variance. ROIs specific to each direction were selected for analysis. For the analysis of all directions, data for ROIs not present in one or more directions were filled with zeros, though the results were unchanged when instead substituted with random numbers drawn from a normal distribution with a mean of 0 and standard deviation of 0.05. iNM effects transfer to swallow and baseline-tongue at rest blocks during which there is no neuromodulation applied.

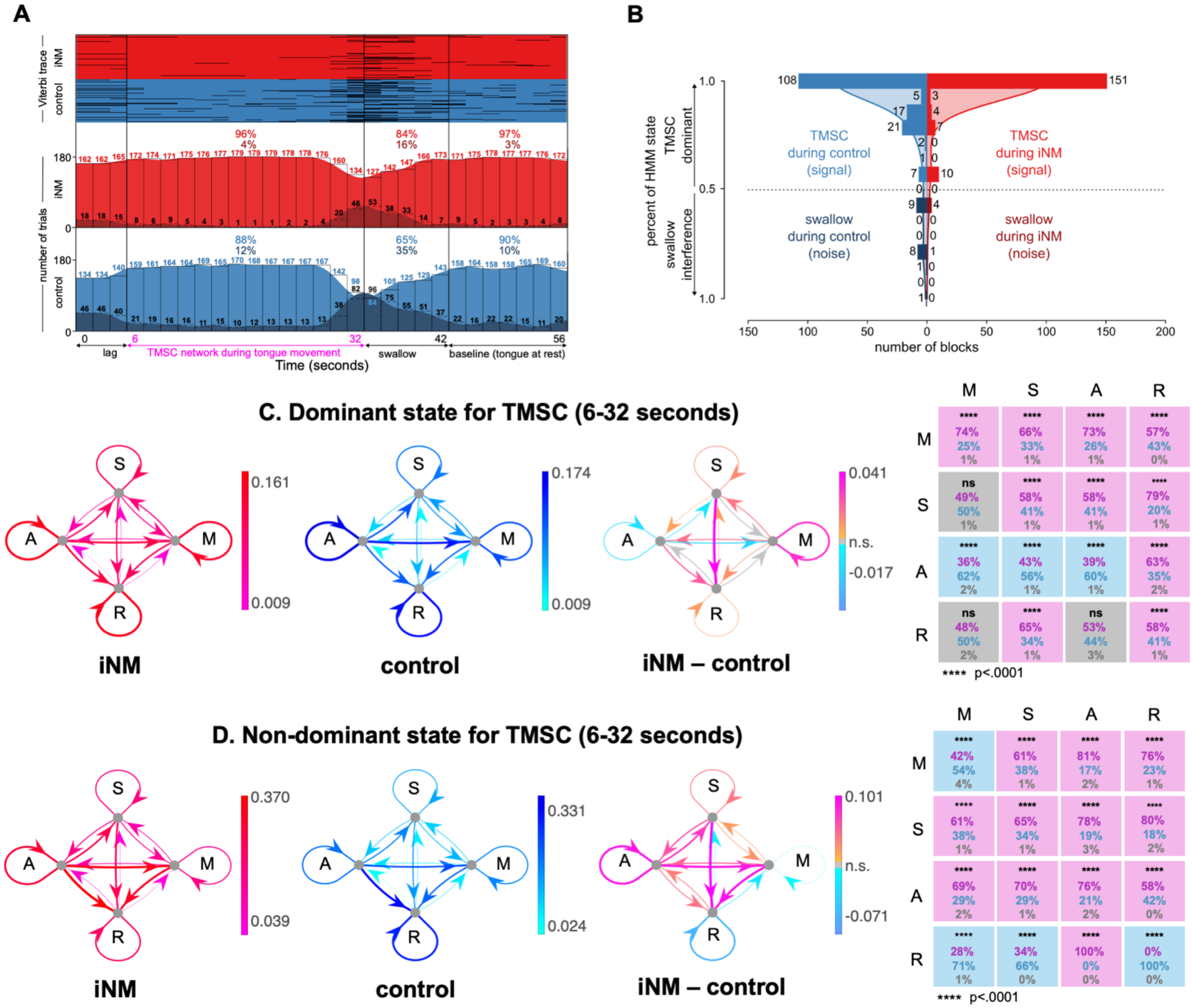


**Extended Data Figure 9: Effect of iNM on network interactions across Left tongue motor control directions elucidated via dynamic causal modeling: A.** Our autoregressive hidden Markov model (ARHMM) was fit to the brain areas that selected for Left tongue movement direction (6-32 seconds). Model inference was then performed on the entire timeseries (0-56 seconds). (top) Viterbi trace for iNM (red) and control (blue) trials that characterize each brain state during Left TM, swallow, and baseline intervals for each condition. (bottom) Histogram of the number of trials corresponding to Left TMSC (light red/blue) or swallow (dark red/blue) brain state for each time repetition, divided into the iNM (red) and control (blue) conditions. Percentages correspond to the fraction of trials within each brain state across the Left TMSC, swallow, or baseline intervals. **B.** The histogram compares the signal (Left TMSC state, red/blue) and noise (swallow state, dark red/dark blue) by displaying the number of trials in each bin that correspond to each of the two states (total trials (iNM vs. control): 151 vs. 108 (Left TMSC state), 5 vs. 19 (swallow state). Shaded curves represent the kernel density estimate (KDE) fit to the binned data for the iNM and control conditions. The KDE was calculated by the summation of normal distributions centered at each data point, followed by normalization, to generate a smoothed estimate of the histogram bins. KDE bandwidth was estimated via Scott's method, and boundary corrections at 0 and 1 employed reflection of the data across the boundaries. **C-D.** For the dominant (Left TMSC) or non-dominant state, the strength of ROI-to-ROI connections within networks (M = Motor network, S = Sensory network, A = Attention network, R = Reward network) is quantified as the mean absolute value of the autoregressive coefficients during the iNM (fuchsia/red) or control (light/dark blue) condition, or the difference in absolute values between conditions (orange/fuchsia and light/dark blue indicate increased/decreased during iNM, respectively). At far right, significant increases (fuchsia), decreases (light blue), or insignificant differences (gray) in these network-to-network interactions are shown for iNM compared to control. Percentages indicate the fraction of region-to-region interactions within each network-to-network connection that are increased (fuchsia), decreased (light blue), or insignificant (gray) in iNM compared to control. Correction for multiple comparisons used an FDR of 0.05.

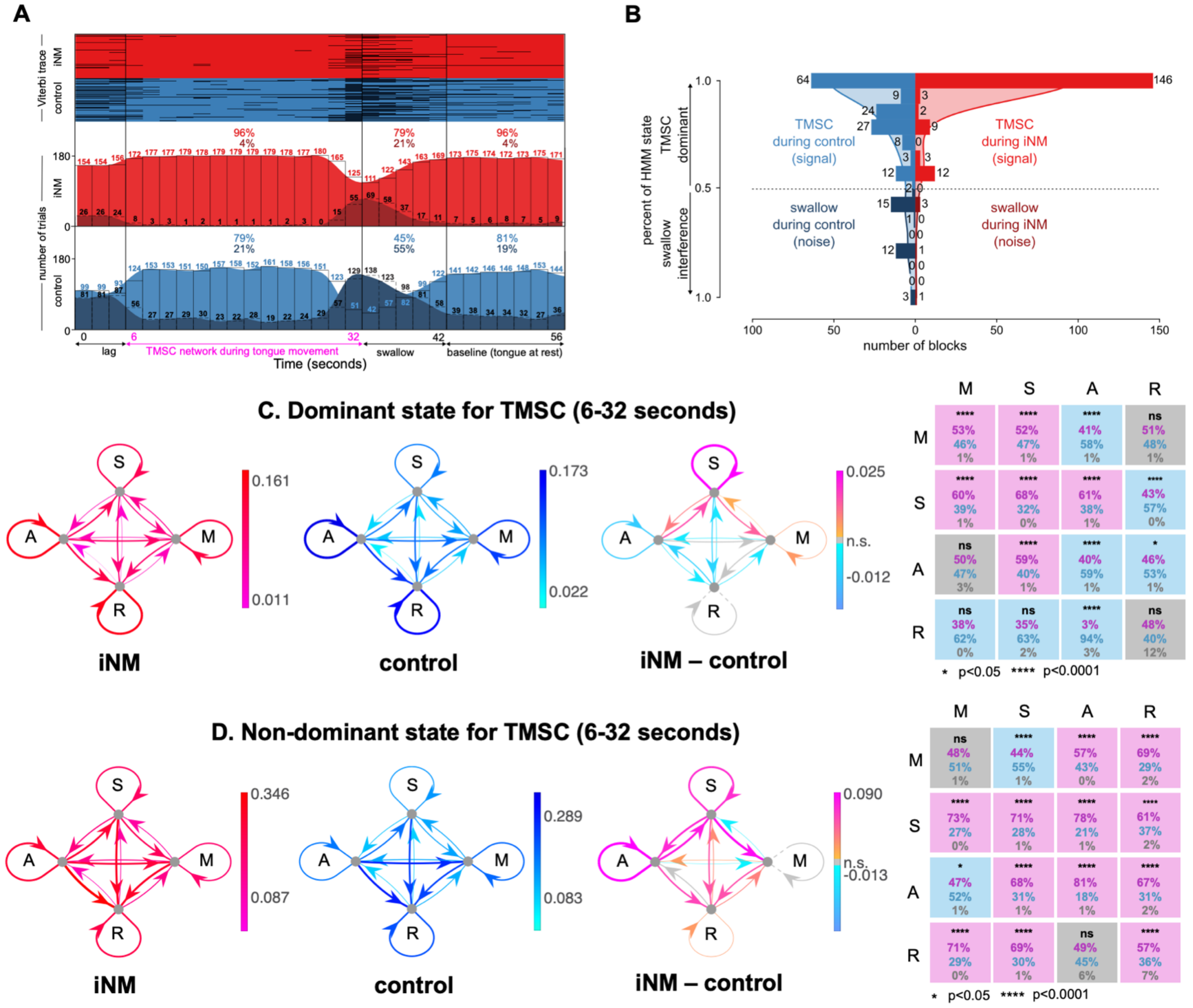


**Extended Data Figure 10: Effect of iNM on network interactions across Right tongue motor control directions elucidated via dynamic causal modeling: A.** Our autoregressive hidden Markov model (ARHMM) was fit to the brain areas that selected for Right tongue movement direction (6-32 seconds). Model inference was then performed on the entire timeseries (0-56 seconds). (top) Viterbi trace for iNM (red) and control (blue) trials that characterize each brain state during Right TM, swallow, and baseline intervals for each condition. (bottom) Histogram of the number of trials corresponding to Right TMSC (light red/blue) or swallow (dark red/blue) brain state for each time repetition, divided into the iNM (red) and control (blue) conditions. Percentages correspond to the fraction of trials within each brain state across the Right TMSC, swallow, or baseline intervals. **B.** The histogram compares the signal (Right TMSC state, red/blue) and noise (swallow state, dark red/dark blue) by displaying the number of trials in each bin that correspond to each of the two states (total trials (iNM vs. control): 146 vs. 64 (Right TMSC state), 5 vs. 33 (swallow state)). Shaded curves represent the kernel density estimate (KDE) fit to the binned data for the iNM and control conditions. The KDE was calculated by the summation of normal distributions centered at each data point, followed by normalization, to generate a smoothed estimate of the histogram bins. KDE bandwidth was estimated via Scott's method, and boundary corrections at 0 and 1 employed reflection of the data across the boundaries. **C-D.** For the dominant (Right TMSC) or non-dominant state, the strength of ROI-to-ROI connections within networks (M = Motor network, S = Sensory network, A = Attention network, R = Reward network) is quantified as the mean absolute value of the autoregressive coefficients during the iNM (fuchsia/red) or control (light/dark blue) condition, or the difference in absolute values between conditions (orange/fuchsia and light/dark blue indicate increased/decreased during iNM, respectively). At far right, significant increases (fuchsia), decreases (light blue), or insignificant differences (gray) in these network-to-network interactions are shown for iNM compared to control. Percentages indicate the fraction of region-to-region interactions within each network-to-network connection that are increased (fuchsia), decreased (light blue), or insignificant (gray) in iNM compared to control. Correction for multiple comparisons used an FDR of 0.05.

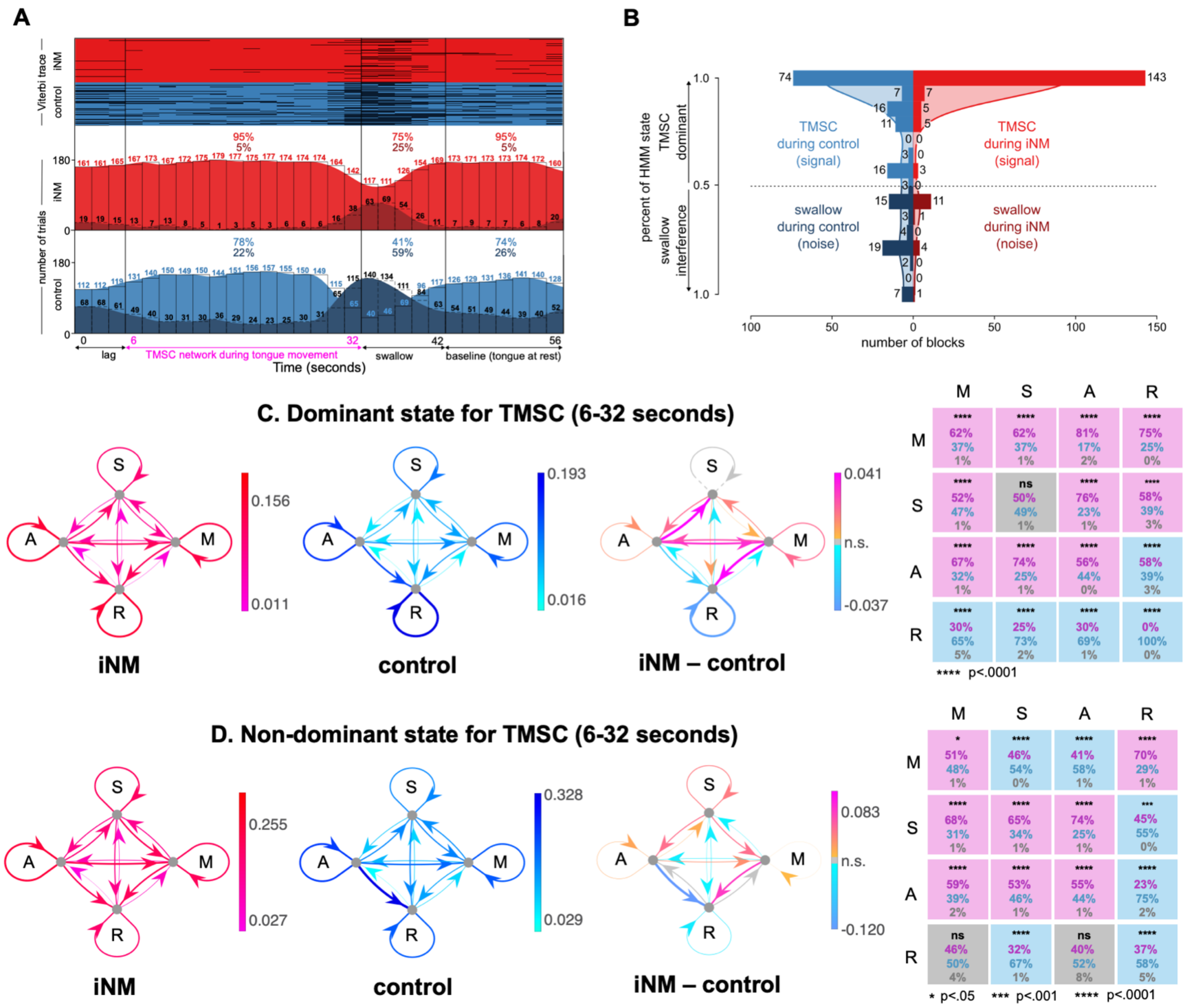


**Extended Data Figure 11: Effect of iNM on network interactions across Up tongue motor control directions elucidated via dynamic causal modeling: A.** Our autoregressive hidden Markov model (ARHMM) was fit to the brain areas that selected for Up tongue movement direction (6-32 seconds). Model inference was then performed on the entire timeseries (0-56 seconds). (top) Viterbi trace for iNM (red) and control (blue) trials that characterize each brain state during Up TM, swallow, and baseline intervals for each condition. (bottom) Histogram of the number of trials corresponding to Up TMSC (light red/blue) or swallow (dark red/blue) brain state for each time repetition, divided into the iNM (red) and control (blue) conditions. Percentages correspond to the fraction of trials within each brain state across the Up TMSC, swallow, or baseline intervals. **B.** The histogram compares the signal (Up TMSC state, red/blue) and noise (swallow state, dark red/dark blue) by displaying the number of trials in each bin that correspond to each of the two states (total trials (iNM vs. control): 143 vs. 74 (Up TMSC state), 17 vs. 53 (swallow state)). Shaded curves represent the kernel density estimate (KDE) fit to the binned data for the iNM and control conditions. The KDE was calculated by the summation of normal distributions centered at each data point, followed by normalization, to generate a smoothed estimate of the histogram bins. KDE bandwidth was estimated via Scott's method, and boundary corrections at 0 and 1 employed reflection of the data across the boundaries. **C-D.** For the dominant (Up TMSC) or non-dominant state, the strength of ROI-to-ROI connections within networks (M = Motor network, S = Sensory network, A = Attention network, R = Reward network) is quantified as the mean absolute value of the autoregressive coefficients during the iNM (fuchsia/red) or control (light/dark blue) condition, or the difference in absolute values between conditions (orange/fuchsia and light/dark blue indicate increased/decreased during iNM, respectively). At far right, significant increases (fuchsia), decreases (light blue), or insignificant differences (gray) in these network-to-network interactions are shown for iNM compared to control. Percentages indicate the fraction of region-to-region interactions within each network-to-network connection that are increased (fuchsia), decreased (light blue), or insignificant (gray) in iNM compared to control. Correction for multiple comparisons used an FDR of 0.05.

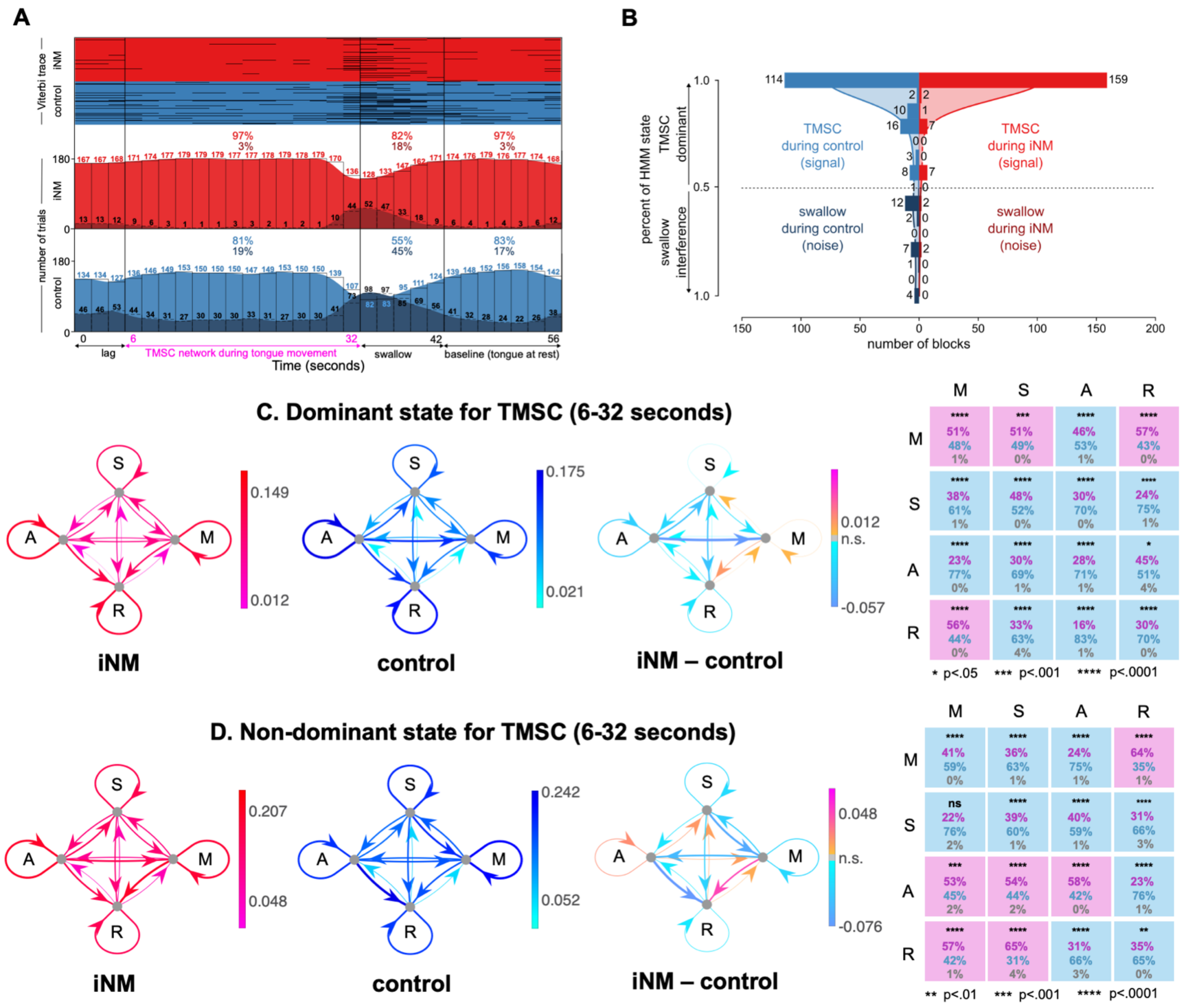


**Extended Data Figure 12: Effect of iNM on network interactions across Down tongue motor control directions elucidated via dynamic causal modeling: A.** Our autoregressive hidden Markov model (ARHMM) was fit to the brain areas that selected for Down tongue movement direction (6-32 seconds). Model inference was then performed on the entire timeseries (0-56 seconds). (top) Viterbi trace for iNM (red) and control (blue) trials that characterize each brain state during Down TM, swallow, and baseline intervals for each condition. (bottom) Histogram of the number of trials corresponding to Down TMSC (light red/blue) or swallow (dark red/blue) brain state for each time repetition, divided into the iNM (red) and control (blue) conditions. Percentages correspond to the fraction of trials within each brain state across the Down TMSC, swallow, or baseline intervals. **B.** The histogram compares the signal (Down TMSC state, red/blue) and noise (swallow state, dark red/dark blue) by displaying the number of trials in each bin that correspond to each of the two states (total trials (iNM vs. control): 159 vs. 114 (Down TMSC state), 4 vs. 26 (swallow state). Shaded curves represent the kernel density estimate (KDE) fit to the binned data for the iNM and control conditions. The KDE was calculated by the summation of normal distributions centered at each data point, followed by normalization, to generate a smoothed estimate of the histogram bins. KDE bandwidth was estimated via Scott’s method, and boundary corrections at 0 and 1 employed reflection of the data across the boundaries. **C-D.** For the dominant (Down TMSC) or non-dominant state, the strength of ROI-to-ROI connections within networks (M = Motor network, S = Sensory network, A = Attention network, R = Reward network) is quantified as the mean absolute value of the autoregressive coefficients during the iNM (fuchsia/red) or control (light/dark blue) condition, or the difference in absolute values between conditions (orange/fuchsia and light/dark blue indicate increased/decreased during iNM, respectively). At far right, significant increases (fuchsia), decreases (light blue), or insignificant differences (gray) in these network-to-network interactions are shown for iNM compared to control. Percentages indicate the fraction of region-to-region interactions within each network-to-network connection that are increased (fuchsia), decreased (light blue), or insignificant (gray) in iNM compared to control. Correction for multiple comparisons used an FDR of 0.05.

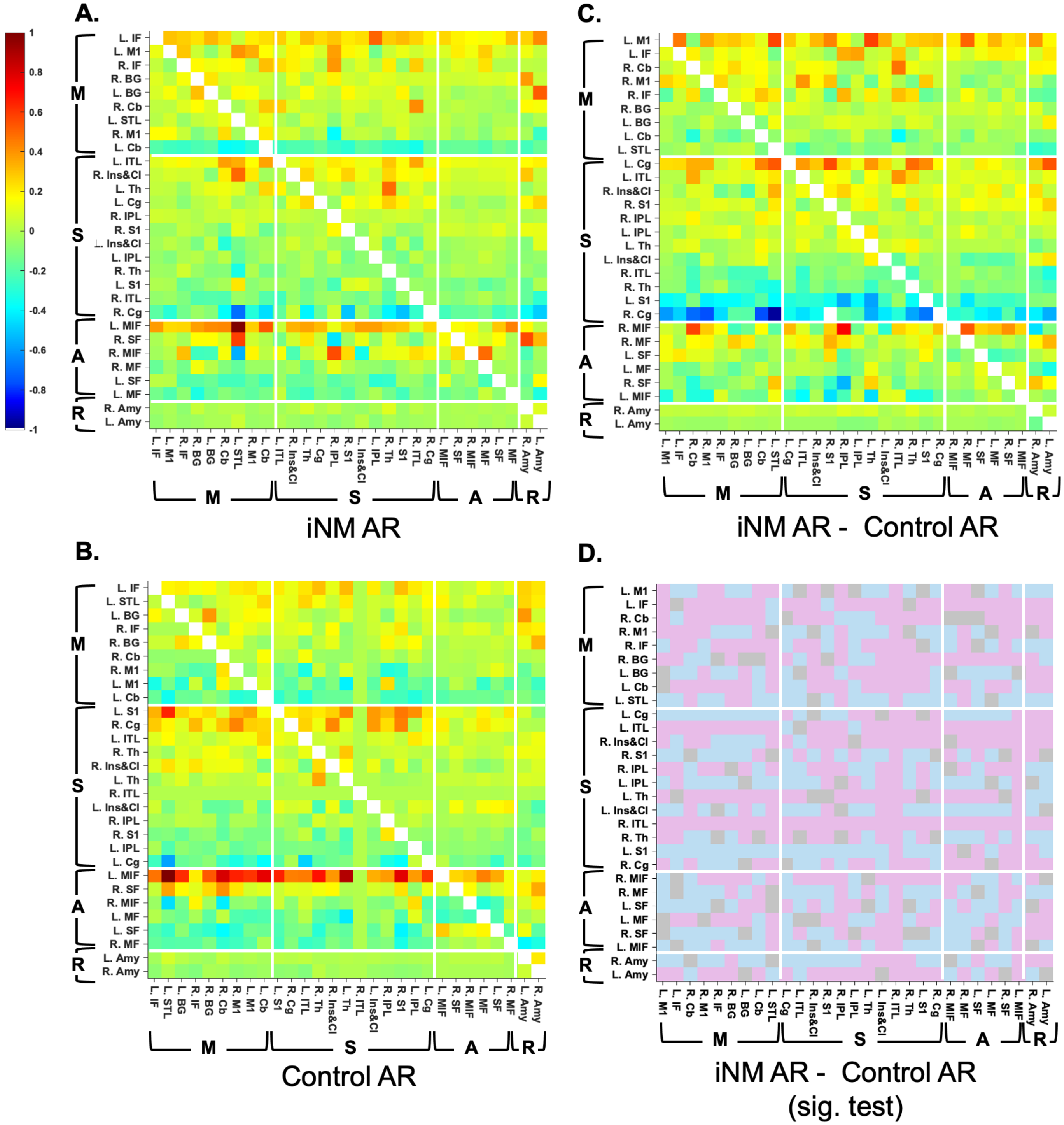


**Extended Data Figure 13: Autoregressive coefficients for the hidden Markov model (HMM) reflecting trials that correspond to tongue movement in the left direction.** The autoregressive component of the HMM correspond to a coefficient matrix that reflected ROI-to-ROI connectivity in data under either the **(A)** iNM and **(B)** control conditions. The heatmaps show the values normalized within the range [-1, +1]. Within each network (M=motor, S=sensory, A=attention, R=reward), ROIs were sorted in descending order according to the average value within each row. The direction of connectivity is from row to column. **(C)** The difference in autoregressive coefficients between the iNM and control conditions is shown. **(D)** The significance of the difference in coefficients was calculated by applying a t-test across the paired data points for each ROI-to-ROI connection (30 subjects; pink, iNM > control; blue, iNM < control; gray, insignificant).

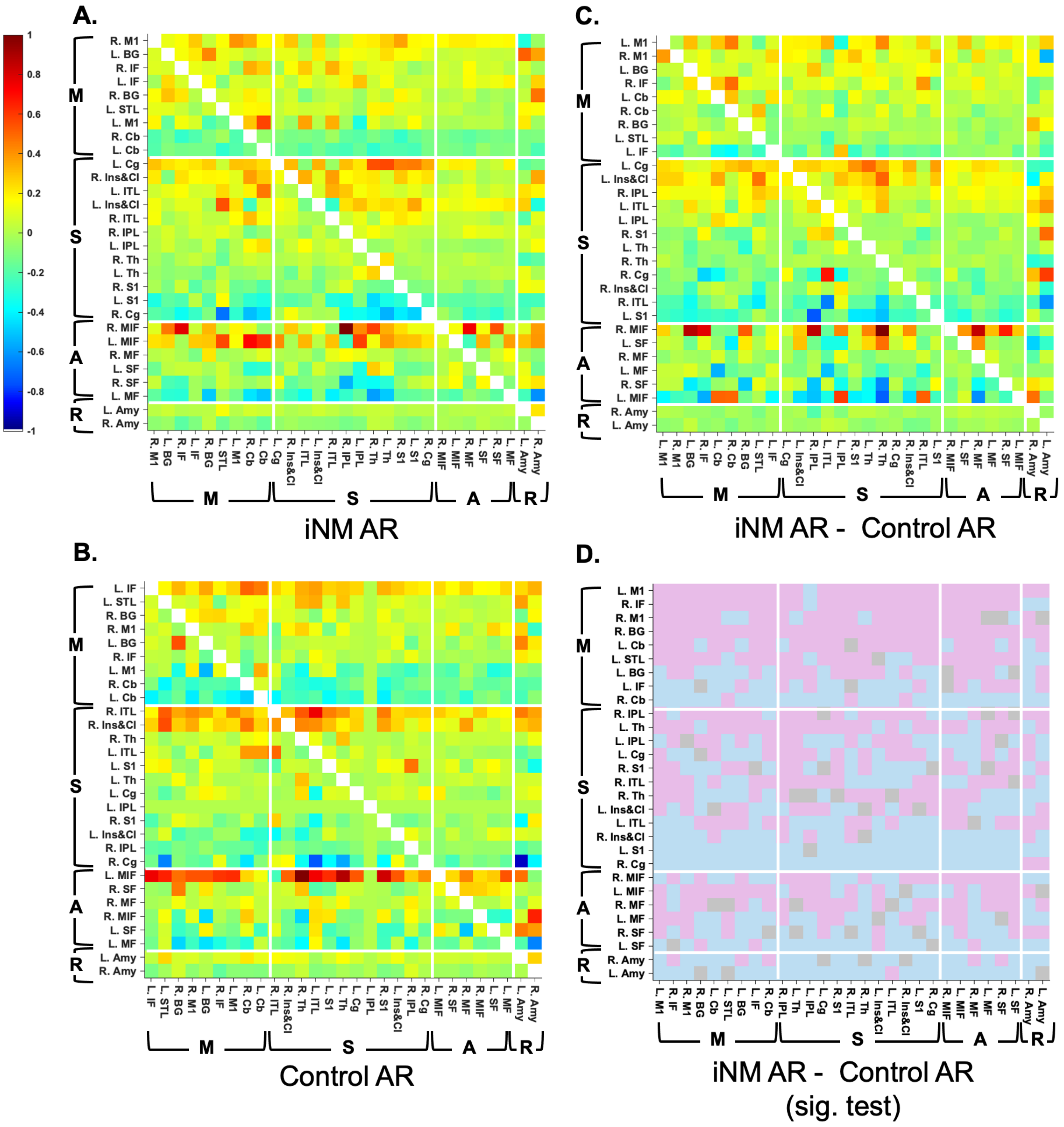


**Extended Data Figure 14: Autoregressive coefficients for the hidden Markov model (HMM) reflecting trials that correspond to tongue movement in the right direction.** Panels **(A-D)** as per caption for Extended Data Figure 12.

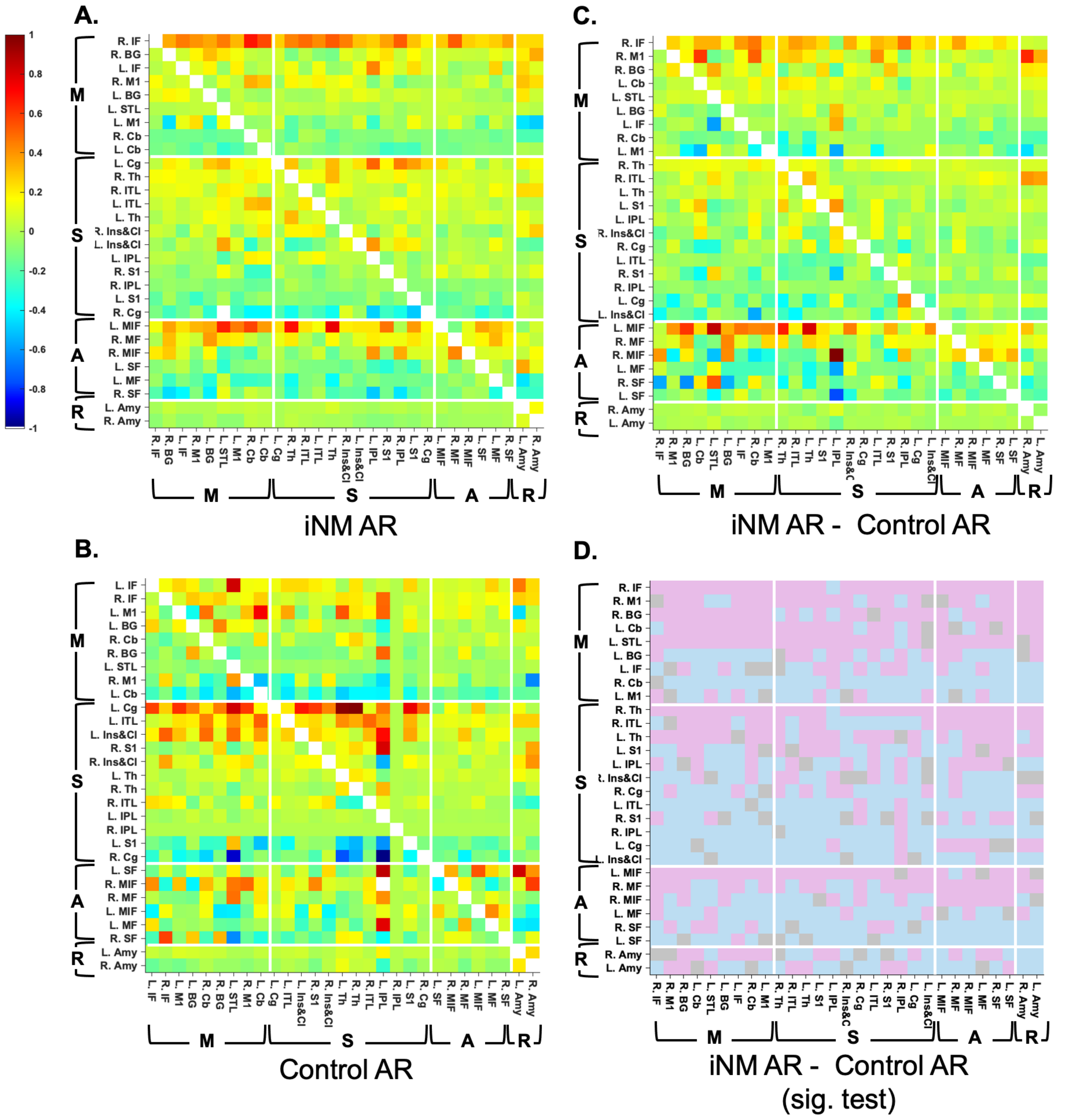


**Extended Data Figure 15: Autoregressive coefficients for the hidden Markov model (HMM) reflecting trials that correspond to tongue movement in the up direction.** Panels **(A-D)** as per caption for Extended Data Figure 12.

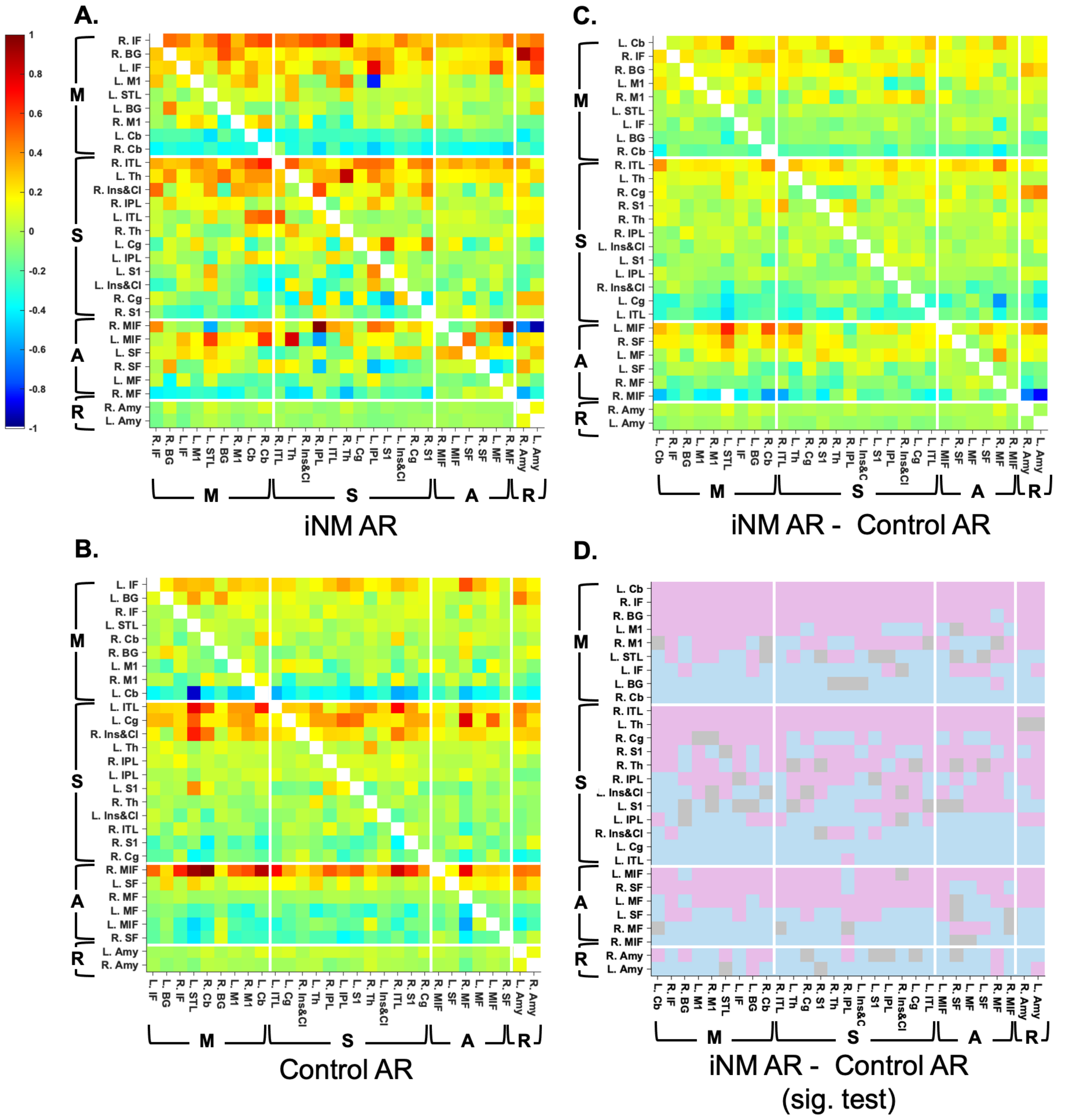


**Extended Data Figure 16: Autoregressive coefficients for the hidden Markov model (HMM) reflecting trials that correspond to tongue movement in the down direction.** Panels **(A-D)** as per caption for Extended Data Figure 12.

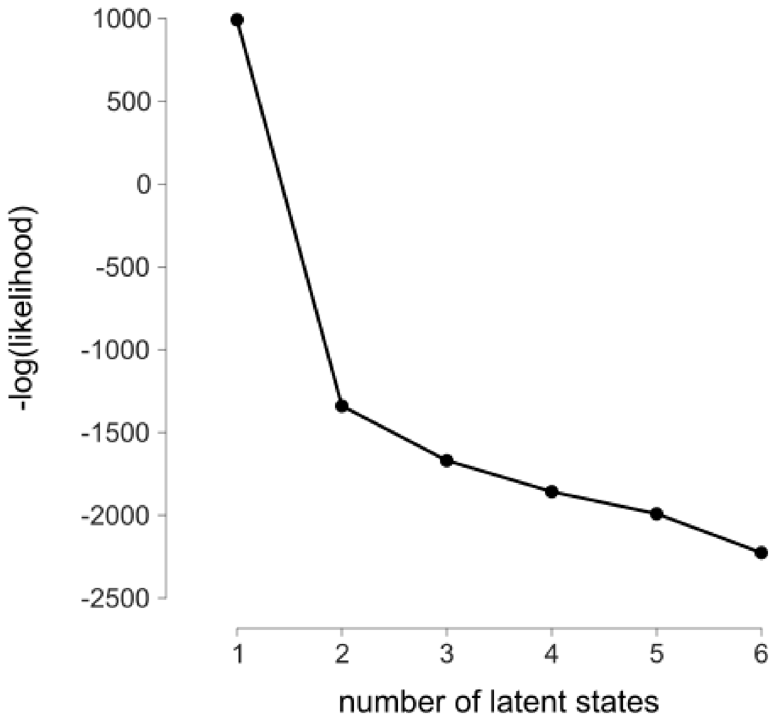


**Extended Data Figure 17**: **Loss function for the autoregressive hidden Markov model (ARHMM) comparing number of latent states.** The ARHMM was fit to the tongue movement phase (6-32 seconds) of the combined dataset, in which trials from all directions (left, right, up, down) and conditions (control, iNM), were merged into a single matrix. Using leave-one-subject-out cross-validation, the model was fit to the tongue movement phase for 29 of 30 subjects, and then tested on the entire timeseries (0-56 seconds) for the remaining subject, repeated across all 30 subjects. The log-likelihood of the test data was calculated and averaged across the 30 subjects for varying number of latent states. The averaged negative log-likelihood, which is reflective of the model loss function, shows an "elbow" at two latent states, suggesting that it best captures the underlying structure of the data without overfitting parameters.

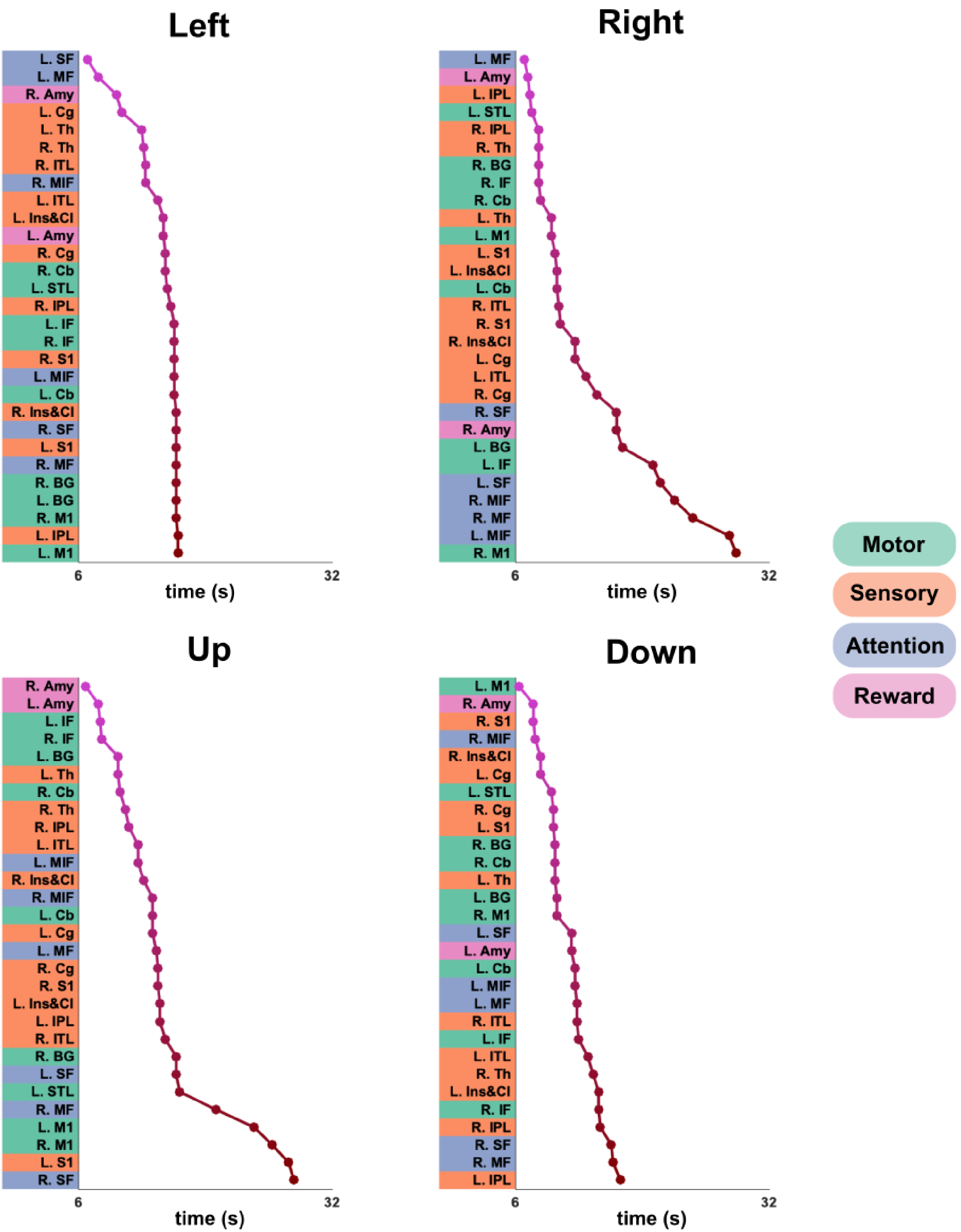


**Extended Data Figure 18**: **Time of earliest peak ROI activity as quantified by the d' score for individual tongue movement directions.** The first peak time of ROI activity during the tongue movement task is shown for each tongue movement direction. ROIs are shown in descending order of peak time and color-coded according to their network (motor, green; sensory, orange; attention, blue; reward, pink). Peak times were calculated by first generating a smoothed estimate of the d' score with cubic interpolation, followed by determining peak times for each ROI using the MATLAB built-in function "peakfinder". The first peak of each ROI was then tabulated.

**Table S1a.** iNM condition: peaks of activated regions for Left TMSC direction selectivity.

| TT_Daemon Atlas Region | BA | Cluster | Cluster Size (# of Voxels) | Region Size (# of Voxels) | RAI Peak Coordinates (mm) | | | Peak Z-score |
|---|---|---|---|---|---|---|---|---|
| | | | | | x | y | z | |
| L precentral (premotor & 1ary motor cortex) | 4, 6 | 1 | 9824 | 588 | 46.5 | 7.5 | 26.5 | 7.7 |
| R precentral (premotor & 1ary motor cortex) | 4, 6 | 1 | 9824 | 502 | -46.5 | 10.5 | 29.5 | 7.3 |
| L postcentral (1ary somatosensory) | 3 | 1 | 9824 | 212 | 58.5 | 13.5 | 26.5 | 7.0 |
| R postcentral (1ary somatosensory) | 3 | | | 259 | -58.5 | 7.5 | 20.5 | 7.2 |
| L thalamus | | 1 | 9824 | 79 | 10.5 | 16.5 | -0.5 | 5.4 |
| R thalamus | | 1 | 9824 | 88 | -10.5 | 22.5 | -9.5 | 6.4 |
| L lentiform & putamen | | 1 | 9824 | 129 | 25.5 | 1.5 | -3.5 | 7.1 |
| R lentiform & putamen | | 1 | 9824 | 158 | -25.5 | 1.5 | -6.5 | 7.3 |
| L insula & claustrum | | 1 | 9824 | 246 | 31.5 | -13.5 | 11.5 | 6.3 |
| R insula & claustrum | | 1 | 9824 | 312 | -34.5 | 7.5 | 14.5 | 6.5 |
| L inferior frontal | 9 | 1 | 9824 | 254 | 34.5 | -10.5 | 23.5 | 5.0 |
| R inferior frontal (operculum) | 45 | 1 | 9824 | 246 | -31.5 | -25.5 | 8.5 | 6.7 |
| L middle frontal | 10 | 1 | 9824 | 217 | 31.5 | -43.5 | 11.5 | 4.3 |
| R middle frontal | 10 | 1 | 9824 | 411 | -31.5 | -40.5 | 23.5 | 5.1 |
| L medial frontal | 24 | 1 | 9824 | 329 | 7.5 | 4.5 | 47.5 | 5.3 |
| R medial frontal (premotor cortex) | 6 | 1 | 9824 | 404 | -4.5 | 7.5 | 53.5 | 5.6 |
| L superior frontal | 10 | 1 | 9824 | 198 | 16.5 | -43.5 | 23.5 | 4.3 |
| R superior frontal (anterior) | 9, 32 | 1 | 9824 | 422 | -13.5 | -19.5 | 44.5 | 6.5 |
| L inferior parietal | 40 | 1 | 9824 | 199 | 49.5 | 40.5 | 41.5 | 4.3 |
| R inferior parietal | 40 | 1 | 9824 | 239 | -46.5 | 49.5 | 47.5 | 4.6 |
| R superior parietal | | 1 | 9824 | 37 | -25.5 | 70.5 | 20.5 | 3.8 |
| L nodule (lobule IX) | | 2 | 829 | 6 | 13.5 | 55.5 | -24.5 | 5.7 |
| L declive (lobule VI – crus 1) | | 2 | 829 | 190 | 31.5 | 61.5 | -21.5 | 5.5 |
| L uvula (lobule VII) | | 2 | 829 | 29 | 34.5 | 67.5 | -24.5 | 4.6 |
| R culmen (lobule VI) | | 1 | 9824 | 65 | -16.5 | 55.5 | -21.5 | 4.8 |
| L inferior temporal | 37 | 2 | 829 | 87 | 46.5 | 61.5 | -15.5 | 6.3 |
| R inferior temporal | | 1 | 9824 | 80 | -43.5 | 58.5 | -12.5 | 5.7 |
| L cingulate (middle) | 24, 32 | 1 | 9824 | 278 | 7.5 | 4.5 | 47.5 | 5.3 |
| R cingulate (middle) | | 1 | 9824 | 313 | -4.5 | 7.5 | 32.5 | 3.6 |
| L anterior cingulate | 24, 32 | 1 | 9824 | 37 | 16.5 | -34.5 | 11.5 | 4.1 |
| R anterior cingulate | | 1 | 9824 | 113 | -13.5 | -34.5 | 14.5 | 4.8 |
| R superior temporal | 38 | 1 | 9824 | 86 | -37.5 | -7.5 | -21.5 | 3.6 |
| R substantia nigra | | 1 | 9824 | 11 | -10.5 | 19.5 | -9.5 | 5.8 |
| L amygdala | 28 | 1 | 9824 | 8 | 25.5 | 1.5 | -9.5 | 5.1 |
| R amygdala | 28 | 1 | 9824 | 47 | -25.5 | 1.5 | -9.5 | 6.7 |

**Table S1b.** Control-No-iNM condition: peaks of activated regions for Left TMSC direction selectivity.

| TT_Daemon Atlas Region | BA | Cluster | Cluster Size (# of Voxels) | Region Size (# of Voxels) | RAI Peak Coordinates (mm) | | | Peak Z-score |
|---|---|---|---|---|---|---|---|---|
| | | | | | x | y | z | |
| L precentral (premotor & 1ary motor cortex) | 6, 4 | 1 | 6211 | 504 | 49.5 | 7.5 | 26.5 | 7.9 |
| R precentral (premotor & 1ary motor cortex) | 6, 4 | 1 | 6211 | 408 | -58.5 | 4.5 | 29.5 | 7.7 |
| L postcentral (1ary somatosensory) | 3/1/2 | 1 | 6211 | 174 | 43.5 | 16.5 | 29.5 | 5.8 |
| R postcentral (1ary somatosensory) | 3/1/2 | 1 | 6211 | 218 | -58.5 | 13.5 | 29.5 | 6.1 |
| L thalamus (subthalamic nucleus) | | 4 | 56 | 29 | 10.5 | 16.5 | -0.5 | 5.8 |
| R thalamus (subthalamic nucleus) | | 1 | 6211 | 36 | -13.5 | 16.5 | -0.5 | 6.1 |
| L lentiform, putamen | | 1 | 6211 | 136 | 25.5 | 1.5 | -6.5 | 6.9 |
| R lentiform, putamen | | 1 | 6211 | 165 | -25.5 | 1.5 | -6.5 | 7.1 |
| L insula & claustrum | | 1 | 6211 | 230 | 34.5 | 7.5 | 14.5 | 6.5 |
| R insula & claustrum | | 1 | 6211 | 254 | -34.5 | 7.5 | 14.5 | 6.9 |
| L middle frontal | 6, 8 | 1 | 6211 | 146 | 31.5 | -13.5 | 44.5 | 4.4 |
| R middle frontal | 10 | 1 | 6211 | 117 | -31.5 | -43.5 | 2.5 | 4.3 |
| L medial frontal (supplementary motor area) | 8, 9 | 1 | 6211 | 296 | 7.5 | -25.5 | 47.5 | 5.3 |
| R medial frontal | 9 | 1 | 6211 | 323 | -22.5 | -34.5 | 23.5 | 5.3 |
| L superior medial frontal | 9 | 1 | 6211 | 225 | 10.5 | -46.5 | 35.5 | 4.1 |
| R superior posterior frontal | 10 | 1 | 6211 | 311 | -19.5 | -49.5 | 23.5 | 4.7 |
| L culmen (lobule VI) | | 2 | 259 | 24 | 13.5 | 55.5 | -24.5 | 5.8 |
| R culmen (lobule VI) | | 6 | 40 | 6 | -16.5 | 58.5 | -21.5 | 4.5 |
| L cingulate (caudal) | 24 | 1 | 6211 | 211 | 7.5 | 4.5 | 47.5 | 5.8 |
| R cingulate (caudal) | 24 | 1 | 6211 | 210 | -10.5 | -7.5 | 47.5 | 5.1 |
| L anterior cingulate | | 1 | 6211 | 71 | 16.5 | -19.5 | 20.5 | 5.5 |
| R anterior cingulate | | 1 | 6211 | 80 | -16.5 | -31.5 | 2.5 | 4.6 |
| L superior temporal | 22 | 3 | 162 | 156 | 37.5 | 25.5 | 5.5 | 5.4 |
| R superior temporal | 22 | 1 | 6211 | 195 | -52.5 | 10.5 | 5.5 | 5.1 |
| R parahippocampal (uncus) | | 1 | 6211 | 48 | -31.5 | 4.5 | -24.5 | 3.6 |

**Table S2a.** iNM condition: peaks of activated regions for Right TMSC direction selectivity.

| TT_Daemon Atlas Region | BA | Cluster | Cluster Size (# of Voxels) | Region Size (# of Voxels) | RAI Peak Coordinates (mm) | | | Peak Z-score |
|---|---|---|---|---|---|---|---|---|
| | | | | | x | y | z | |
| L precentral (premotor & 1ary motor) | 6 | 1 | 9467 | 566 | 46.5 | 7.5 | 26.5 | 7.5 |
| R precentral (premotor & 1ary motor) | 6 | 1 | 9467 | 484 | -46.5 | 10.5 | 26.5 | 7.2 |
| L postcentral (1ary somatosensory) | 3 | 1 | 9467 | 196 | 55.5 | 10.5 | 44.5 | 3.8 |
| R postcentral (1ary somatosensory) | 3 | 1 | 9467 | 261 | -58.5 | 10.5 | 20.5 | 6.6 |
| L thalamus | | 1 | 9467 | 105 | 10.5 | 16.5 | 2.5 | 5.9 |
| R thalamus | | 1 | 9467 | 101 | -10.5 | 16.5 | -0.5 | 6.0 |
| L lentiform & putamen | | 1 | 9467 | 165 | 25.5 | 1.5 | -3.5 | 6.7 |
| R lentiform & putamen | | 1 | 9467 | 126 | -25.5 | 1.5 | -6.5 | 6.4 |
| L insula & claustrum | | 1 | 9467 | 305 | 34.5 | -1.5 | 5.5 | 5.5 |
| R insula & claustrum | | 1 | 9467 | 239 | -34.5 | 7.5 | 14.5 | 6.1 |
| L inferior frontal (operculum) | 44, 45 | 1 | 9467 | 244 | 31.5 | -22.5 | 11.5 | 6.5 |
| R inferior frontal | 45 | 1 | 9467 | 260 | -31.5 | -22.5 | 8.5 | 6.2 |
| L middle frontal | 10, 46 | 1 | 9467 | 168 | 31.5 | -46.5 | 20.5 | 4.6 |
| R middle frontal | 9, 10, 46 | 1 | 9467 | 331 | -34.5 | -31.5 | 26.5 | 5.2 |
| L medial frontal (supplementary motor area) | 6 | 1 | 9467 | 294 | 7.5 | 4.5 | 50.5 | 5.7 |
| R medial frontal (supplementary motor area) | 6 | 1 | 9467 | 331 | -4.5 | 7.5 | 53.5 | 5.6 |
| L superior frontal | 10 | 1 | 9467 | 169 | 25.5 | -37.5 | 5.5 | 4.2 |
| R superior frontal | 8, 9 | 1 | 9467 | 387 | -7.5 | -31.5 | 50.5 | 5.7 |
| L nodule & dentate (lobule IX) | | 2 | 537 | 29 | 13.5 | 55.5 | -24.5 | 4.7 |
| L culmen (lobule VI) | | 2 | 537 | 50 | 34.5 | 55.5 | -18.5 | 4.4 |
| R culmen & dentate (lobule IV-V) | | 1 | 9467 | 145 | -16.5 | 58.5 | -21.5 | 5.7 |
| R declive | | 1 | 9467 | 258 | -28.5 | 82.5 | -15.5 | 5.1 |
| L inferior parietal lobule | 40 | 1 | 9467 | 56 | 43.5 | 16.5 | 29.5 | 6.0 |
| R inferior parietal lobule | 40 | 1 | 9467 | 297 | -49.5 | 43.5 | 44.5 | 4.9 |
| R superior parietal lobule | 7 | 1 | 9467 | 32 | -22.5 | 67.5 | 41.5 | 4.2 |
| L inferior temporal (fusiform) | 37 | 2 | 537 | 76 | 43.5 | 58.5 | -15.5 | 5.6 |
| R inferior temporal (fusiform) | | 1 | 9467 | 107 | -49.5 | 55.5 | -0.5 | 3.9 |
| L middle cingulate | 24 | 1 | 9467 | 268 | 10.5 | -10.5 | 35.5 | 4.8 |
| R middle cingulate | 24, 32 | 1 | 9467 | 316 | -10.5 | -16.5 | 38.5 | 4.7 |
| L superior temporal | 28, 36 | 1 | 9467 | 208 | 37.5 | -13.5 | -21.5 | 4.0 |
| R middle & superior temporal | 22 | 1 | 9467 | 91 | -46.5 | 25.5 | -3.5 | 4.0 |
| L substantia nigra | | 1 | 9467 | 11 | 13.5 | 19.5 | -3.5 | 4.6 |
| R substantia nigra | | 1 | 9467 | 9 | -11.5 | 19.5 | -6.5 | 4.5 |
| L superior temporal lobe (uncus) | 28, 36 | 1 | 9467 | 20 | 28.5 | 4.5 | -24.5 | 4.0 |
| L amygdala | | 1 | 9467 | 14 | 25.5 | 1.5 | -9.5 | 5.8 |
| R amygdala | | 1 | 9467 | 9 | -25.5 | 1.5 | -9.5 | 5.4 |

**Table S2b.** Control-No iNM: peaks of activated regions for Right TMSC direction selectivity

| TT_Daemon Atlas Region | BA | Cluster | Cluster Size (# of Voxels) | Region Size (# of Voxels) | RAI Peak Coordinates (mm) | | | Peak |
|---|---|---|---|---|---|---|---|---|
| | | | | | x | y | z | Z-score |
| L precentral (premotor & 1ary motor) | 4, 6 | 1 | 6169 | 507 | 46.5 | 7.5 | 26.5 | 7.8 |
| R precentral (premotor & 1ary motor) | 4, 6 | 1 | 6169 | 380 | -58.5 | 4.5 | 29.5 | 7.3 |
| L postcentral (1ary somatosensory) | 3/1/2 | 1 | 6169 | 137 | -49.5 | 10.5 | 23.5 | 7.0 |
| R postcentral (1ary somatosensory) | 3/1/2 | 1 | 6169 | 197 | -61.5 | 7.5 | 17.5 | 6.9 |
| L thalamus (subthalamic nucleus) | | 7 | 34 | 3 | 10.5 | 16.5 | -0.5 | 5.8 |
| R thalamus (subthalamic nucleus) | | 8 | 21 | 2 | -13.5 | 16.5 | -0.5 | 5.1 |
| L lentiform & putamen | | 1 | 6169 | 129 | 25.5 | 1.5 | -6.5 | 7.2 |
| R lentiform & putamen | | 1 | 6169 | 86 | -25.5 | 1.5 | -6.5 | 7.0 |
| L insula & claustrum | | 1 | 6169 | 189 | 34.5 | 7.5 | 14.5 | 5.2 |
| R insula & claustrum | | 1 | 6169 | 214 | -34.5 | 7.5 | 14.5 | 5.9 |
| L inferior frontal (operculum) | 9 | 1 | 6169 | 80 | 43.5 | -10.5 | 26.5 | 3.8 |
| L middle frontal | 9, 46 | 1 | 6169 | 181 | 28.5 | -19.5 | 20.5 | 5.7 |
| R middle frontal | 10, 11 | 1 | 6169 | 103 | -34.5 | -49.5 | 2.5 | 3.4 |
| L medial frontal | 9 | 1 | 6169 | 262 | 22.5 | -28.5 | 26.5 | 5.2 |
| R medial frontal | 9 | 1 | 6169 | 343 | -16.5 | -34.5 | 29.5 | 5.0 |
| L superior frontal | 10, 12 | 1 | 6169 | 262 | 28.5 | -43.5 | 8.5 | 4.7 |
| R superior frontal | 9 | 1 | 6169 | 293 | -16.5 | -46.5 | 26.5 | 5.7 |
| L nodule & dentate (lobule IX) | | 4 | 49 | 37 | 13.5 | 55.5 | -24.5 | 5.9 |
| R culmen (lobule VI) | | 2 | 292 | 40 | -16.5 | 58.5 | -21.5 | 5.5 |
| R tuber (lobule VIIB) | | 2 | 292 | 3 | -49.5 | 67.5 | -24.5 | 3.9 |
| L inferior temporal (fusiform) | 37 | 3 | 129 | 37 | 43.5 | 58.5 | -15.5 | 3.4 |
| R inferior temporal (fusiform) | 20, 37 | 2 | 292 | 34 | -37.5 | 58.5 | -12.5 | 4.1 |
| L middle cingulate (supplementary motor area) | 24, 32 | 1 | 6169 | 272 | 4.5 | 4.5 | 47.5 | 5.6 |
| R cingulate (supplementary motor area) | 24 | 1 | 6169 | 324 | -7.5 | 4.5 | 47.5 | 5.1 |
| L anterior cingulate | | 1 | 6169 | 66 | 10.5 | -31.5 | 11.5 | 4.3 |
| L superior temporal | 38 | 1 | 6169 | 213 | 34.5 | -4.5 | -21.5 | 4.2 |
| R superior temporal | 41, 42 | 1 | 6169 | 76 | -43.5 | 28.5 | 8.5 | 3.8 |
| L amygdala | 28 | 1 | 6169 | 29 | 25.5 | 1.5 | -9.5 | 6.1 |
| R amygdala | 28 | 1 | 6169 | 17 | -25.5 | 1.5 | -9.5 | 6.4 |

**Table S3a.** iNM condition: peaks of activated regions for Up TMSC direction selectivity.

| TT_Daemon Atlas Region | BA | Cluster | Cluster Size (# of Voxels) | Region Size (# of Voxels) | RAI Peak Coordinates (mm) | | | Peak Z-score |
|---|---|---|---|---|---|---|---|---|
| | | | | | x | y | z | |
| L precentral (premotor & 1ary motor cortex) | 4 | 1 | 8400 | 538 | 58.5 | 10.5 | 26.5 | 7.8 |
| R precentral (premotor & 1ary motor cortex) | 6 | 1 | 8400 | 514 | -49.5 | 7.5 | 29.5 | 7.2 |
| L postcentral (caudal – 1ary somatosensory) | 3, 2, 1 | 1 | 8400 | 203 | 55.5 | 25.5 | 35.5 | 4.4 |
| R postcentral (rostral – 1ary somatosensory) | 3a | 1 | 8400 | 272 | -58.5 | 7.5 | 20.5 | 7.2 |
| L thalamus | | 4 | 32 | 15 | 10.5 | 16.5 | -0.5 | 4.2 |
| R thalamus | | 6 | 17 | 12 | -10.5 | 16.5 | -0.5 | 4.8 |
| L lentiform & putamen | | 3 | 131 | 80 | 25.5 | 1.5 | -6.5 | 6.1 |
| R lentiform & putamen | | 1 | 8400 | 113 | -28.5 | 4.5 | -6.5 | 6.2 |
| L insula | | 1 | 8400 | 166 | 31.5 | -25.5 | 11.5 | 5.7 |
| R insula | | 1 | 8400 | 215 | -40.5 | 4.5 | 17.5 | 5.7 |
| L inferior frontal | 9 | 1 | 8400 | 208 | 55.5 | -7.5 | 26.5 | 7.0 |
| R inferior frontal | 10 | 1 | 8400 | 311 | -31.5 | -22.5 | 8.5 | 6.5 |
| L middle frontal | 9 | 1 | 8400 | 156 | 25.5 | -22.5 | 26.5 | 4.6 |
| R middle frontal | 10 | 1 | 8400 | 396 | -31.5 | -37.5 | 17.5 | 5.3 |
| L medial frontal | 9 | 1 | 8400 | 217 | 25.5 | -34.5 | 20.5 | 5.3 |
| R medial frontal | 6 | 1 | 8400 | 376 | -1.5 | 7.5 | 53.5 | 5.1 |
| L superior frontal | 8, 9 | 1 | 8400 | 75 | 10.5 | -28.5 | 44.5 | 4.6 |
| R superior frontal | 9 | 1 | 8400 | 290 | -10.5 | -46.5 | 32.5 | 4.6 |
| L culmen (lobule IV-V crus 1) | | 2 | 833 | 101 | 34.5 | 49.5 | -18.5 | 5.3 |
| L declive (lobule VI) | | 2 | 833 | 183 | 31.5 | 58.5 | -18.5 | 5.1 |
| L nodule (lobule IX) | | 2 | 833 | 3 | 13.5 | 55.5 | -24.5 | 4.7 |
| R declive (lobule VI) | | 1 | 8400 | 238 | -34.5 | 67.5 | -21.5 | 5.5 |
| R culmen (lobule IV-V crus 1) | | 1 | 8400 | 146 | -34.5 | 40.5 | -21.5 | 4.9 |
| L inferior parietal | 40 | 1 | 8400 | 138 | 52.5 | 37.5 | 50.5 | 4.2 |
| R inferior parietal | 40 | 1 | 8400 | 252 | -52.5 | 37.5 | 44.5 | 4.5 |
| L inferior temporal (fusiform) | 19, 37 | 2 | 833 | 109 | 46.5 | 64.5 | -15.5 | 6.4 |
| R inferior temporal (fusiform) | 37, 20 | 1 | 8400 | 149 | -43.5 | 55.5 | -12.5 | 6.9 |
| L cingulate (middle) | 32 | 1 | 8400 | 205 | 13.5 | -4.5 | 44.5 | 4.5 |
| R cingulate | 32 | 1 | 8400 | 205 | -22.5 | -10.5 | 29.5 | 4.6 |
| R anterior cingulate | | 1 | 8400 | 107 | -22.5 | -25.5 | 26.5 | 5.4 |
| L superior temporal | 22 | 1 | 8400 | 50 | 61.5 | 1.5 | 8.5 | 4.7 |
| R superior temporal | 22 | 1 | 8400 | 33 | -61.5 | 4.5 | 8.5 | 5.1 |
| L substantia nigra | | 4 | 32 | 6 | 10.5 | 19.5 | -9.5 | 3.8 |

**Table S3b.** Control-No-iNM: Peaks of activated regions for Up TMSC direction selectivity.

| TT_Daemon Atlas Region | BA | Cluster | Cluster Size (# of Voxels) | Region Size (# of Voxels) | RAI Peak Coordinates (mm) | | | Peak Z-score |
|---|---|---|---|---|---|---|---|---|
| | | | | | x | y | z | |
| L precentral (premotor & 1ary motor cortex) | 4, 6 | 3 | 791 | 406 | 55.5 | 4.5 | 20.5 | 8.0 |
| R precentral (premotor & 1ary motor cortex) | 4, 6 | 4 | 776 | 342 | -61.5 | 4.5 | 20.5 | 7.3 |
| L postcentral (1ary somatosensory) | 3, 1, 2 | 3 | 791 | 153 | 64.5 | 13.5 | 29.5 | 6.0 |
| R postcentral (1ary somatosensory) | 3, 1, 2 | 4 | 776 | 191 | -49.5 | 13.5 | 53.5 | 3.8 |
| L thalamus | | 12 | 11 | 5 | 10.5 | 16.5 | -0.5 | 4.5 |
| R thalamus | | 11 | 12 | 9 | -13.5 | 16.5 | -0.5 | 5.0 |
| L lentiform & putamen | | 8 | 127 | 61 | 25.5 | 1.5 | -6.5 | 7.2 |
| R lentiform & putamen | | 7 | 140 | 49 | -25.5 | 1.5 | -6.5 | 6.7 |
| L insula & claustrum | | 2 | 821 | 78 | 31.5 | -22.5 | 17.5 | 4.5 |
| R insula & claustrum | | 4 | 776 | 99 | -34.5 | 7.5 | 14.5 | 6.0 |
| L inferior frontal | 10 | 2 | 821 | 70 | 43.5 | -40.5 | 2.5 | 3.2 |
| R inferior frontal | 47 | 1 | 1383 | 74 | -31.5 | -13.5 | -12.5 | 4.2 |
| L middle frontal | 8 | 2 | 821 | 118 | 25.5 | -10.5 | 35.5 | 4.3 |
| R middle frontal | 46 | 1 | 1383 | 104 | -28.5 | -43.5 | 11.5 | 5.2 |
| L medial frontal | 10, 46 | 2 | 821 | 167 | 19.5 | -46.5 | 11.5 | 5.1 |
| R medial frontal | 10 | 1 | 1383 | 232 | -22.5 | -37.5 | 17.5 | 4.7 |
| L superior frontal | 6, 8 | 2 | 821 | 142 | 16.5 | -16.5 | 50.5 | 4.6 |
| R superior frontal | 8 | 1 | 1383 | 156 | -10.5 | -25.5 | 44.5 | 4.6 |
| L declive (superior vermian lobule VI) | | 6 | 356 | 81 | 43.5 | 61.5 | -18.5 | 5.4 |
| L culmen (lobule IV-V) | | 6 | 356 | 37 | 31.5 | 52.5 | -15.5 | 4.5 |
| R declive (lobule VI) | | 5 | 447 | 120 | -34.5 | 58.5 | -15.5 | 5.4 |
| R culmen (lobule IV-V) | | 5 | 447 | 64 | -31.5 | 40.5 | -18.5 | 3.7 |
| L inferior temporal (fusiform) | | 2 | 833 | 63 | 43.5 | 58.5 | -15.5 | 5.2 |
| R inferior temporal (fusiform) | | 1 | 8400 | 49 | -40.5 | 49.5 | -15.5 | 4.4 |
| L cingulate (middle) | 32, 6 | 1 | 1383 | 120 | 7.5 | 4.5 | 47.5 | 5.4 |
| R cingulate (middle) | 32 | 1 | 1383 | 170 | -16.5 | -16.5 | 32.5 | 5.1 |
| L superior temporal | | 3 | 791 | 27 | 58.5 | 1.5 | 8.5 | 4.4 |
| R superior temporal | | 4 | 776 | 68 | -61.5 | 7.5 | 8.5 | 4.6 |
| L amygdala | | 8 | 127 | 14 | 25.5 | 1.5 | -9.5 | 6.3 |
| R amygdala | | 7 | 140 | 23 | -25.5 | 1.5 | -9.5 | 6.2 |

**Table S4a.** iNM condition: Peaks of activated regions for Down TMSC direction selectivity

| TT_Daemon Atlas Region | BA | Cluster | Cluster Size (# of Voxels) | Region Size (# of Voxels) | RAI Peak Coordinates (mm) | | | Peak Z-score |
|---|---|---|---|---|---|---|---|---|
| | | | | | x | y | z | |
| L precentral (premotor & 1ary motor cortex) | 4, 6 | 1 | 5582 | 508 | 58.5 | 10.5 | 26.5 | 8.0 |
| R precentral (premotor & 1ary motor cortex) | 4, 6 | 1 | 5582 | 427 | -49.5 | -4.5 | 8.5 | 4.9 |
| L postcentral (1ary somatosensory) | 40 | 1 | 5582 | 198 | 58.5 | 28.5 | 20.5 | 3.5 |
| R postcentral (1ary somatosensory) | 3/1/2 | 1 | 5582 | 263 | -58.5 | 13.5 | 29.5 | 7.4 |
| L thalamus | | 4 | 190 | 42 | 10.5 | 16.5 | -0.5 | 5.5 |
| R thalamus | | 4 | 190 | 56 | -10.5 | 16.5 | -0.5 | 6.2 |
| L lentiform | | 1 | 5582 | 127 | 25.5 | 4.5 | -3.5 | 6.8 |
| R lentiform | | 1 | 5582 | 109 | -25.5 | 1.5 | -6.5 | 6.2 |
| L insula & claustrum | | 1 | 5582 | 218 | 31.5 | -16.5 | 5.5 | 6.8 |
| R insula & claustrum | | 1 | 5582 | 194 | -31.5 | -19.5 | 8.5 | 5.7 |
| L inferior frontal (operculum) | 44 | 1 | 5582 | 218 | 55.5 | -4.5 | 23.5 | 6.6 |
| R inferior frontal (orbital) | 47 | 1 | 5582 | 260 | -52.5 | -16.5 | -0.5 | 4.1 |
| L middle frontal | 10 | 1 | 5582 | 83 | 28.5 | -37.5 | 11.5 | 4.7 |
| R middle frontal | 10 | 1 | 5582 | 180 | -43.5 | -49.5 | 11.5 | 3.0 |
| L medial frontal | 6 | 1 | 5582 | 168 | 7.5 | 4.5 | 50.5 | 6.0 |
| R medial frontal | 6 | 1 | 5582 | 268 | -4.5 | 7.5 | 53.5 | 5.7 |
| L superior frontal | 10 | 1 | 5582 | 84 | 31.5 | -43.5 | 14.5 | 4.5 |
| R superior frontal | 9 | 1 | 5582 | 180 | -16.5 | -46.5 | 23.5 | 4.6 |
| L uvula (crus 1 - lobule VII) | | 3 | 845 | 33 | 28.5 | 64.5 | -24.5 | 4.7 |
| L nodule & ventral dentate (lobule IX) | | 3 | 845 | 27 | 13.5 | 55.5 | -24.5 | 4.2 |
| L culmen (crus 1 - lobule VI) | | 3 | 845 | 99 | 43.5 | 43.5 | -27.5 | 3.7 |
| R culmen & ventral dentate (crus 1 - lobule VI) | | 2 | 1076 | 119 | -19.5 | 58.5 | -21.5 | 4.6 |
| L inferior parietal | 40 | 1 | 5582 | 71 | 49.5 | 52.5 | 47.5 | 3.7 |
| R inferior parietal | 40 | 1 | 5582 | 253 | -37.5 | 49.5 | 41.5 | 4.7 |
| R superior parietal | 7 | 1 | 5582 | 23 | -28.5 | 52.5 | 38.5 | 4.6 |
| L cingulate (middle) | 24, 32 | 1 | 5582 | 144 | 7.5 | -10.5 | 38.5 | 4.3 |
| R cingulate (middle) | 32 | 1 | 5582 | 178 | -7.5 | -16.5 | 38.5 | 5.0 |
| L anterior cingulate | | 1 | 5582 | 34 | 13.5 | -31.5 | 23.5 | 3.6 |
| R anterior cingulate | 33 | 1 | 5582 | 46 | -4.5 | -10.5 | 23.5 | 4.3 |
| L superior temporal | | 1 | 5582 | 100 | 37.5 | 1.5 | -21.5 | 3.5 |

**Table S4b.** Control-No-iNM: Peaks of activated regions for Down TMSC direction selectivity

| TT_Daemon Atlas Region | BA | Cluster | Cluster Size (# of Voxels) | Region Size (# of Voxels) | RAI Peak Coordinates (mm) | | | Peak Z-score |
|---|---|---|---|---|---|---|---|---|
| | | | | | x | y | z | |
| L precentral (premotor & 1ary motor cortex) | 3a | 1 | 909 | 420 | 52.5 | 4.5 | 23.5 | 7.7 |
| R precentral (premotor & 1ary motor cortex) | 3a | 2 | 791 | 314 | -61.5 | 4.5 | 26.5 | 7.3 |
| L postcentral (1ary somatosensory) | 3/1/2 | 1 | 909 | 149 | 58.5 | 13.5 | 26.5 | 6.7 |
| R postcentral (1ary somatosensory) | 3/1/2 | 2 | 791 | 186 | -58.5 | 13.5 | 29.5 | 7.0 |
| L thalamus (subthalamic nucleus) | | 13 | 12 | 8 | 10.5 | 16.5 | -0.5 | 4.7 |
| R thalamus (subthalamic nucleus) | | 14 | 10 | 8 | -13.5 | 16.5 | -0.5 | 4.5 |
| L lentiform & putamen | | 8 | 114 | 58 | 25.5 | 1.5 | -6.5 | 6.8 |
| R lentiform & putamen | | 7 | 139 | 69 | -25.5 | 1.5 | -6.5 | 7.2 |
| L insula & claustrum | | 1 | 909 | 146 | 34.5 | 7.5 | 14.5 | 5.2 |
| R insula & claustrum | | 2 | 791 | 157 | -37.5 | 4.5 | 14.5 | 5.4 |
| L inferior frontal (operculum) | 47 | 12 | 17 | 100 | 37.5 | -31.5 | -6.5 | 4.1 |
| L middle frontal | 6, 8 | 11 | 42 | 47 | 19.5 | -19.5 | 53.5 | 4.3 |
| R middle frontal | 9 | 10 | 51 | 1 | -25.5 | -28.5 | 23.5 | 3.7 |
| L medial frontal | 9, 10 | 6 | 276 | 98 | 22.5 | -31.5 | 23.5 | 4.5 |
| R medial frontal | 9, 10 | 10 | 51 | 161 | -22.5 | -37.5 | 14.5 | 4.0 |
| L superior frontal | 10 | 6 | 276 | 91 | 31.5 | -43.5 | 14.5 | 4.6 |
| R superior frontal (posterior) | 9 | 3 | 461 | 67 | -16.5 | -49.5 | 23.5 | 4.4 |
| L uvula (lobule VII – crus 2) | | 5 | 387 | 10 | 31.5 | 79.5 | -24.5 | 3.4 |
| R declive (lobule VI – crus 1) | | 4 | 431 | 136 | -22.5 | 61.5 | -18.5 | 5.2 |
| R declive (lobule VI – crus 1) | | 4 | 431 | 52 | -22.5 | 55.5 | -15.5 | 5.0 |
| L middle cingulate | 24 | 3 | 461 | 65 | 10.5 | 4.5 | 47.5 | 4.7 |
| R middle cingulate | 24 | 3 | 461 | 93 | -10.5 | 4.5 | 47.5 | 5.0 |
